# New loci and candidate genes in spring two-rowed barley detected through meta-analysis of a field trial European network

**DOI:** 10.1101/2024.09.04.611234

**Authors:** Francesc Montardit-Tarda, Ana M Casas, William TB Thomas, Florian Schnaithmann, Rajiv Sharma, Salar Shaaf, Chiara Campoli, Joanne Russell, Luke Ramsay, Stefano Delbono, Marko Jääskeläinen, Maitry Paul, Frederick L Stoddard, Andrea Visioni, Andrew J Flavell, Klaus Pillen, Benjamin Kilian, Andreas Graner, Laura Rossini, Robbie Waugh, Luigi Cattivelli, Alan H Schulman, Alessandro Tondelli, Ernesto Igartua

**Affiliations:** Estación Experimental de Aula Dei – Consejo Superior de Investigaciones Científicas (EEAD-CSIC), Avenida Montañana 1005, 500059, Zaragoza, España; James Hutton Institute, Errol Road, Invergowrie, Dundee DD2 5DA, United Kingdom; Martin-Luther-Univ. Halle-Wittenberg, Betty-Heimann-Str. 3, 06120 Halle (Saale), Germany; Leibniz Institute of Plant Genetics and Crop Plant Reasearch (IPK), Corrensstrasse 3, 06466 Gatersleben, Germany; Scotland’s Rural College, Peter Wilson Building, The King’s Buildings, West Mains Road, Edinburgh EH9 3JG, United Kingdom; Università degli Studi di Milano, Via Celoria 2, 20133 Milano, Italy; Council for Agricultural Research and Economics (CREA), Research Centre for Genomics and Bionformatics, Via San Protaso 302, 29017 Fiorenzuola d’Arda, Italy; Institute of Biotechnology, University of Helsinki, FI-00014 Helsinki, Finland; Viikki Plant Sciences Centre, University of Helsinki, FI-00014 Helsinki, Finland; Natural Resources Institute Finland (LUKE), FI-00014 Helsinki, Finland; International Center for Agricultural Research in the Dry Areas (ICARDA), Rabat 10100, Morocco; Dundee University at SCRI, Invergowrie, Dundee DD2 5DA, United Kingdom; Global Crop Diversity Trust, Platz Der Vereinten Nationen 7, 53113 Bonn, Germany

**Keywords:** agronomic traits, barley, breeding history, GWAS, meta-analysis, *Vrs1*

## Abstract

This study contributes new knowledge on quantitative trait loci (QTLs) and candidate genes for adaptive traits and yield in two-rowed spring barley. A meta-analysis of a network of field trials, varying in latitude and sowing date, with 151 cultivars across several European countries, increased QTL detection power compared to single-trial analyses. The traits analysed were heading date (HD), plant height (PH), thousand-grain weight (TGW), and grain yield (GY). Breaking down the analysis by the main genotype-by-environment trends revealed QTLs and candidate genes specific to conditions like sowing date and latitude. A historical look on the evolution of QTL frequencies revealed that early selection focused on PH and TGW, likely due to their high heritability. GY selection occurred later, facilitated by reduced variance in other traits. The study observed that favourable alleles for plant height were often fixed before those for grain yield and TGW. Some regions showed linkage in repulsion, suggesting targets for future breeding. Several candidate genes were identified, including known genes and new candidates based on orthology with rice. Remarkably, the *deficiens* allele of gene *Vrs1*, appears associated to higher GY. These findings provide valuable insights for barley breeders aiming to improve yield, and other agronomic traits.

**Highlight:** A dense genome-wide meta-analysis provides new QTLs, reveals breeding history trends and identifies new candidate genes for yield, plant height, grain weight and heading time of spring barley.

## Introduction

Barley (*Hordeum vulgare* L.) is the fourth-ranking cereal in the world, and one of the most important crops in Europe, in terms of cultivation area and economic relevance (Dawson et al., 2015, Looseley et al., 2020). In Europe, barley has been the subject of intensive breeding for over 100 years. Competitive breeding in the spring two-rowed pool, with thorough use of the traditional “cross the best with the best and hope for the best” strategy has increased concerns about possible genetic erosion in the cultivated germplasm pool. This process has led to the preferential selection of some genomic regions, and to an overall decrease in genetic diversity, particularly in the spring barley pool (Kolodinska Brantestam et al., 2004; Dziurdziak et al., 2022; Schmidt et al., 2023). Indeed, Tondelli et al. (2013) detected signs of extinction of diversity in some genomic regions. Intensive breeding activities usually produce, inadvertently or consciously, fixation of alleles with large effects on important target traits. However, genetic variation is still present (Tondelli et al., 2013), although finding QTLs, even with small effects, becomes harder. One way to detect minor QTLs is by relying on extensive phenotyping and meta-analysis (Muñoz-Amatriaín et al., 2020). In many European regions, barley with spring growth habit is sown between February and May, to avoid harsh winters. This is mandatory in Nordic countries and other European areas with harsh winters, particularly in Eastern Europe, but cultivation of spring-type barley occurs throughout Europe. Its relevance is increasing for two reasons. On the one hand, increasing winter temperatures allow its cultivation in areas of Europe (like Germany, Italy, Spain, or Switzerland) where winter barley and autumn/winter sowings were prevalent. On the other hand, the main economic boost for barley breeding in Europe has been, and still is, malting quality, a sector largely dominated by spring two-rowed types, consequently, breeding efforts have been particularly intense within this pool, giving rise to malting cultivars as productive as the best feed barleys.

Genome-wide Association (GWA) studies have been widely used in barley to find genomic regions of interest for a large variety of agronomic characters (Igartua et al., 2019; Thomas, 2020). In some cases, candidate genes were identified and validated, making for straightforward breeding. Even so, this is only possible when large diversity panels are available, combined with enough marker density. Marker density provided by the 50k SNP chip (Bayer et al., 2017) gives the opportunity to search for candidate genes in association studies. In narrow germplasm sets, linkage disequilibrium (LD) should be high, hence relatively low marker density is sufficient to pinpoint QTL regions in GWA studies (GWAS). However, to differentiate cultivars that are very close, and be able to track alleles of candidate genes, higher marker densities eventually are needed. This is likely the case of the cultivated spring two-rowed barley pool. Dense genotyping can be achieved using exome capture data, which is available in barley (Mascher et al., 2013; Russell et al., 2016; Chen et al., 2022). Another aspect which has not been fully exploited in GWA is the information from multi-trial studies. With few exceptions (for instance Bustos-Korts et al., 2019), these are analysed as the mean of all trials, or by identifying the intersection of associated markers between single-trials analyses. These methods do not make full use of the potential of independent effects tested in multiple sites to detect QTLs (Muñoz-Amatriain et al., 2020).

This study aims at finding new QTLs for relevant agronomic traits in spring two-rowed barley cultivars, and at indicating new candidate genes that may become new targets for barley breeding in Europe.

## Materials and methods

### Plant material

A collection of 164 spring two-rowed cultivars was tested in the framework of the international projects EXBARDIV (http://pgrc.ipk-gatersleben.de/barleynet/projects_exbardiv.php) and CLIMBAR (https://project-wheel.faccejpi.net/climbar/). The two projects included extensive sets of genotypes, but only spring two-rowed cultivars common to both projects were kept for this study. A principal component analysis for marker data was carried out with the *SNPRelate* package (Zheng et al., 2012) in R (R Core Team, 2024). Cultivars clearly outside the principal cloud of points were filtered out (Fig. S1). Discarded cultivars either originated in southern Europe (likely representing distinct germplasm pools) or had introgressions from exotic parents. A total of 151 cultivars that did not show a clear population structure were kept for further analyses (Table S1).

### Phenotypic evaluation, curation and analysis

Field trials were carried out in 2009 and 2010 within the EXBARDIV project, and in 2016 and 2017 in the framework of the CLIMBAR project, in the United Kingdom, Finland, Germany, Italy, Spain and Morocco (Table 1, Table S2). All trials consisted of plots of four to eight rows, 2-3 m long, and 1-1.5 m wide, in two replicates, following alpha-lattice designs, with plots managed according to local practices for sowing rate and chemical inputs. Flowering time (HD, days from sowing to appearance of the spike out of the flag leaf sheath, Z55, according to Zadoks et al. 1974), plant height (PH, cm, length from the ground to the tip of the spike, without awns, average of five plants), grain yield after combine harvest (GY), and thousand grain weight (TGW) were recorded. Raw phenotypic data from EXBARDIV were retrieved for the 2009 and 2010 seasons, partially reported in Xu et al. (2018). Phenotypic data were curated for outlier data. Two trials (MAR17 and DE2-10) were fully discarded due either to consistently low correlation coefficients with the rest (Fig. S2) or to low overall data quality. Data from 16 trials were kept for HD, PH and GY, while TGW was recorded from 15 trials only.

**Table 1.** Field trial network; locations and years. Phenotype means, standard deviation, and broad sense heritability.

| Country | Trial code | Sowing date | Heading time (DAS) |  | Plant height (cm) |  | Grain yield (t/ha) |  | Thousand grain weight (g) |  |
| --- | --- | --- | --- | --- | --- | --- | --- | --- | --- | --- |
|  |  |  | Mean | SD | Mean | SD | Mean | SD | Mean | SD |
| Morocco | MAR16 | 06/01/16 | 104.0 | 4.5 | 71.3 | 7.2 | 3.25 | 0.55 | NA | NA |
| Spain | ESP16 | 11/11/15 | 163.7 | 6.6 | 63 | 8.8 | 2.98 | 0.67 | 33.2 | 3.3 |
| Spain | ESP17 | 15/11/16 | 154.1 | 4.6 | 54.1 | 6.5 | 2.85 | 0.5 | 39.1 | 3 |
| Italy | ITA16 | 05/11/15 | 167.8 | 4.0 | 86.4 | 6.9 | 7.98 | 1.13 | 38.4 | 5.2 |
| Italy | ITA17 | 08/11/16 | 174.5 | 4.9 | 84.3 | 6.9 | 6.79 | 1.03 | 46.6 | 4.6 |
| Italy | ITA09 | 17/02/09 | 96.3 | 2.9 | 65.3 | 6.1 | 4.55 | 0.74 | 43.6 | 3.1 |
| Italy | ITA10 | 01/03/10 | 93.6 | 3.8 | 52.4 | 5.8 | 3.14 | 0.69 | 42.5 | 2.6 |
| Germany | DE1-09 | 31/03/09 | 62.8 | 3.3 | 76.2 | 7.4 | 3.91 | 0.76 | 35.9 | 4.6 |
| Germany | DE1-10 | 06/04/10 | 67.5 | 2.2 | 83.2 | 8.3 | 4.74 | 0.76 | 46.5 | 3.7 |
| Germany | DE2-09 | 03/04/09 | 66.5 | 2.9 | 89.9 | 12.8 | 6.55 | 0.66 | 50.1 | 3.4 |
| United Kingdom | GBR09 | 25/03/09 | 90.7 | 2.5 | 72.3 | 10.8 | 5.51 | 0.63 | 46.1 | 3.4 |
| United Kingdom | GBR10 | 01/04/10 | 78.9 | 2.6 | 82 | 9.7 | 5.24 | 0.76 | 46.9 | 3.9 |
| United Kingdom | GBR16 | 16/03/16 | 89.5 | 2.3 | 89.8 | 14.2 | 6.21 | 0.73 | 50.4 | 3.2 |
| United Kingdom | GBR17 | 29/03/17 | 83.4 | 2.2 | 88.1 | 10.4 | 7.04 | 0.79 | 51.0 | 3.6 |
| Finland | FIN16 | 11/05/16 | 53.3 | 3.4 | 49.4 | 6.3 | 2.94 | 0.6 | 35.8 | 3.1 |
| Finland | FIN17 | 19/05/17 | 53.9 | 1.9 | 69.9 | 7.5 | 5.43 | 0.56 | 49.2 | 3.6 |
| Heritability ($h^2$) | | | 0.822 | | 0.869 | | 0.216 | | 0.836 | |

Best linear unbiased estimators (BLUEs) were calculated with Genstat 20 (VSN International, 2022). In each trial, the best spatial correction model was used, with the simplest model being a randomized complete block design; the full model including replicates, autoregressive order 1 in rows and columns, and additional contributions from significant random row and column factors (Table S2). Chi-square tests were performed for models differing by a single factor. The most parsimonious model for each trait was chosen, the last in which the inclusion of a spatial correction factor improved the model significantly. If a given cultivar had missing data in three or less trials, its phenotype was imputed with the value corresponding to the percentile of that trait for the missing cultivar in the average of the remaining trials (21, 21, 27 and 17 imputed values for HD, PH, GY and TGW, respectively).

An additive main effects and multiplicative interaction analysis (AMMI) was done for each trait with Genstat 20 (VSN International, 2022). These analyses were used to cluster the trials into mega-environments, following the main direction of genotype by environment interaction (GEI) per trait.

### Genotyping

The lines were genotyped with the 50k Illumina Infinium SNP Array (Bayer et al., 2017). Missing data were imputed with Beagle 5.0 (Browning et al., 2018), as described in Bretani et al. (2022). After imputation, 40639 markers remained. For further analysis, markers with a minor allele frequency equal to or higher than 0.05 were kept (28988 markers). Physical positions of markers were retrieved from both MorexV1 (Mascher et al., 2017) and MorexV3 (Mascher et al., 2021). Additional genotyping for flowering time genes and *Vrs1* (main gene determining spike type) was performed for all lines, with specific markers developed as described in Table S1.

### Genome-Wide Association Analysis and meta-analysis

Association analyses at the single trial level were carried out for all phenotypes and trials using the mixed linear model (MLM) implemented in the GAPIT package (Wang and Zhang, 2021) in R (R Core Team, 2024), with a genomic kinship matrix for adjustment of relatedness, calculated with a randomly selected set of 10% of markers (Fig. S3).

The results of a single trial GWA per trait were meta-analysed with the software METAL (Willer et al., 2010), using the sample size strategy, for the whole set of trials, and for the best grouping of trials indicated by the AMMI analysis. For grain yield, ITA16 and ITA17 were discarded for meta-analysis due to the high dispersion within each trial. A meta-GWA threshold was calculated as the minimum *p*-value detected by 1000 meta-analysis of 1000 permutations per trial, for each phenotype and combination of trials. Markers with a higher -log10(*p*-val) than the threshold were declared as a marker-trait association (MTA). Neighbouring MTAs were grouped into single QTL with two different criteria. First, MTAs from the same chromosome were grouped according to a cluster analysis, as in Looseley et al. (2020). The marker with the largest association per QTL was declared as flag marker. Then, flag markers from the same chromosome, which were in the same LD block (detailed below), were merged.

### Linkage disequilibrium analysis

A basal genomic LD threshold was computed. This threshold was estimated as the square of the 95^th^ percentile of the distribution of unlinked r^2^ values (square root transformed), as in Breseghello and Sorrells (2006). This distribution was fitted with the values of the interchromosomal r^2^ between 200 random markers per chromosome, discounting the population structure using the r2v parameter, using the R package *LDcorSV* (Mangin et al., 2012), which considers kinship relatedness. For each chromosome, intrachromosomic LD block size was calculated using r2v, for pairwise LD values between 500 random markers per chromosome. Chromosomic LD decay was calculated as the point where a loess regression intercepted with the basal genomic LD, using R package *fitdistrplus* (Delignette-Muller and Dutang, 2015); this procedure served to merge the flag markers of several MTAs into a single QTL.

To search for candidate genes, the confidence region for each QTL was calculated. Local LD decay around each flag marker was estimated fitting a loess regression to the pairwise LD values from the closest 400 markers. The confidence region was defined as the distance from the flag marker to the point where the loess curve decreased to the basal genomic LD threshold. When the loess curves did not converge, the chromosomal LD was used instead to declare confidence intervals.

### Meta-analysis multilocus model

The markers found in the meta-analysis may still present some multicollinearity. To reduce it, sequential multivariate analyses of variance were carried out for each trait, with package *car* (Fox and Weisberg, 2019). The flag markers were introduced as independent variables, and the genotypic values for each trial were the dependent variables. Markers were introduced sequentially at each step, keeping in the model the most significant one at each round. The final model summarizes the set of markers most likely having a combined independent effect for all trials, on each trait.

### GWA enrichment with exome capture markers

Markers based on exome capture (EC) (Chen et al., 2022) were used for the refinement of peak regions found with 50k markers. For each QTL, all available markers (50k + EC markers) within its local LD block were retrieved and a single-trial GWA and its meta-analysis were run again. Exome capture markers with higher -log10(*p*-values) than that of the flag marker were considered indicators of a possible candidate gene. Relevant annotations of homologs from other species and gene expression (Milne et al., 2021; Li et al., 2023) in tissues related to the phenotype were considered as additional pointers for possible candidate genes.

Homologues of the candidate genes from *Arabidopsis thaliana*, rice, and wheat were identified with protein BLAST in order to gather functional information using Ensembl Plants (Yates et al., 2022). Only the top homologue with an identity above 80% was considered. Orthologues of *Oryza sativa subsp. japonica* of each candidate gene were identified with Ensembl Plants. FunRiceGenes (Huang et al., 2022) was used to check if a rice orthologue was a trait-related gene.

### Allele frequency shifts over time of cultivar release

The genotypic panel used is representative of the progression of spring barley breeding in Europe. The wide range in the cultivars’ year of release enables tracking of the fate of QTLs in parallel with the history of European spring barley breeding. The cultivars were grouped by year of release into four groups, with the number of cultivars in parentheses: 1920-1959 (n = 18), 1960-1979 (38), 1980-1999 (69) and 2000-(29), respectively. Mean allele frequencies of 250 rolling windows per chromosome were calculated for the four groups of cultivars. MetaQTLs allele frequencies were also calculated for the same four classes, to describe any trends likely due to breeding. Allele frequency shifts of metaQTLs, and genome-wide rolling windows, were determined as the difference between the allele frequencies of the oldest and most recent group of cultivars.

## Results

### Field trial results

The field trials represented a varied range of latitudes, climates, and edaphic and agronomic conditions. Accordingly, grain yields were highly variable, between 2.8 and 8.0 t ha^-1^, with an overall yield of 4.9 t ha^-1^. Autumn- or winter-sown trials in Spain and Morocco showed the lowest yields, while trials autumn-sown in Italy and spring-sown trials at northern latitudes were more productive (Table 1, Fig. S4). The largest variation in days from sowing to heading was mainly caused by sowing date. The trials that were autumn-sown in Italy or Spain, as well as the winter-sown trial in Morocco, experienced longer cycles, followed by the late-winter-sown trials in Italy (ITA09 and ITA10, sown in February and early March), and by all the spring-sown trials. Among the latter, Scottish trials showed longer seasons than German and Finnish trials. Within sowing dates, those differences in cycle lengths were probably caused by different rates of accumulation of growing degree days. Plant height also varied widely between trial means, from 52 to 90 cm, suggesting a large variability in water availability during the stem elongation phase. Thousand-grain weight (TGW) varied between 33 g, for some southern locations, to 51 g in the Scottish trials, indicating highly variable grain filling conditions. Grain yield trial means showed a tight correlation of 0.84 with plant height (a surrogate of biomass) and a moderate correlation with TGW (0.60), indicating the importance of the biomass formation phase, throughout the entire season, and of the conditions prevalent during grain filling for grain yield build-up (Fig. S5).

For all the traits, both the genotypic and genotype by environment (G x E) interaction effects were significant. In general, G x E was more relevant for GY or TGW than for HD or PH but, in all cases, the genotypic sum of squares was much larger. G x E patterns revealed by the AMMI analysis were different for each trait. For grain yield, most trials formed a tight cloud, except those of ITA16 and ITA17, both being trials with high yields having a large effect on overall G x E variance (Table 2, Fig. S6). For heading date (Fig. 1), the first principal component reflected mainly a difference between sowing dates, with autumn-sown (ESP16, ESP17, ITA16, ITA17) or winter-sown trials (MAR16) showing large positive loadings, spring-sown ones (DE1-09, DE1-10, DE2-09, GBR09, GBR10, GBR16, GBR17, FIN16, FIN17) placed opposite to them, and intermediate-sowing dates (ITA09 and ITA10) in a halfway position. ITA09 and ITA10 were, however, classified as spring trials for meta-analysis. The first principal component explained a large proportion of G x E, and was strongly influenced by the differential behaviour of Nordic cultivars Mona and Saana. These cultivars were relatively later in the northernmost environments, and very early in the autumn-sown trials. This was also evidenced by the comparison of the differences between heading date of these cultivars and the overall mean at each trial (Fig. S7). The distribution of trials over the first component for plant height showed a geographic pattern, with most southern environments (Italy, Morocco and Spain) on one side of the first axis, and most northern towards the opposite side (Finland, Germany and the United Kingdom). For TGW, the unique behaviour of ITA16 was the main cause of the very large first principal component.

**Table 2.**
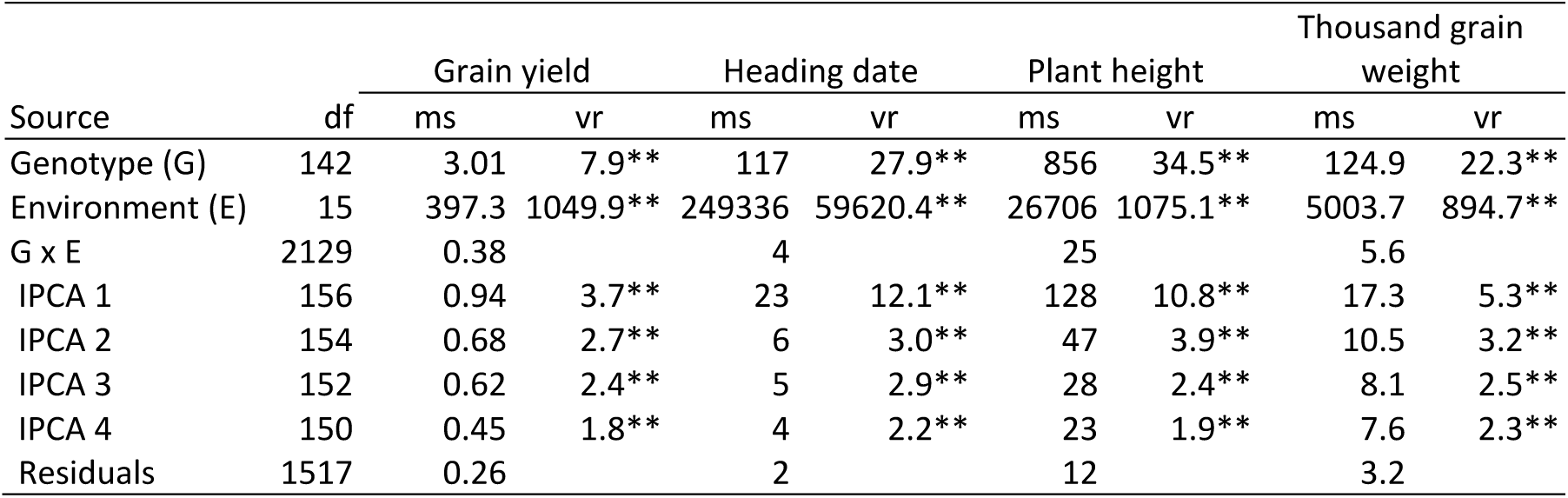
Multi-environment analysis of variance for BLUEs of four traits. Mean squares are presented (ms). As the replicate layer is not present, the variance ratio (vr) for genotype and environment is calculated with the G x E term in the denominator, becoming a stringent test, as this variance adds true G x E variance to the error variance. G x E variance is broken down in the first four components of an AMMI analysis for each trait, each tested for significance against the residual G x E variance left after removing the variance accounted for by each principal component (PC).

**Figure 1.**
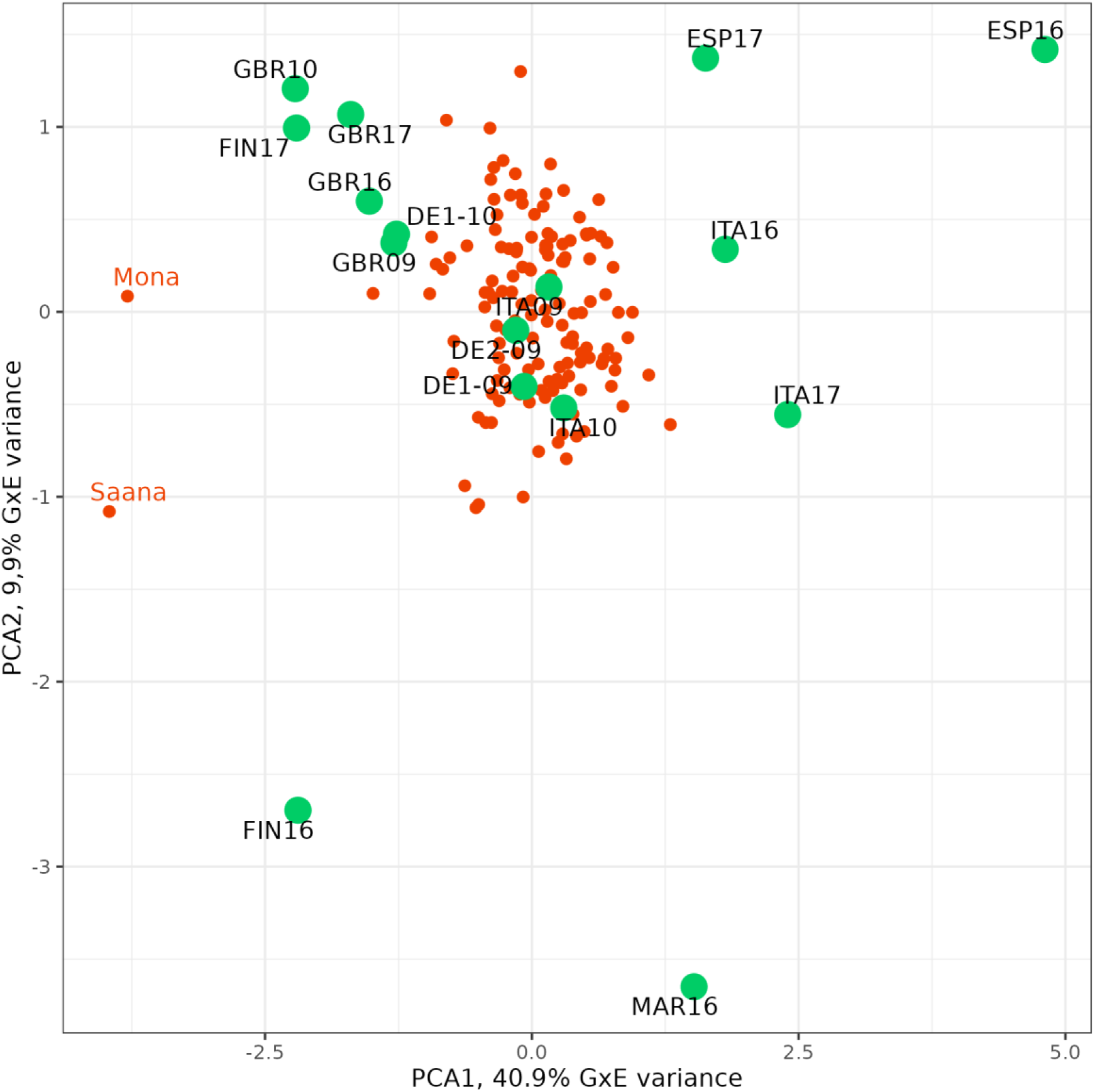
Plot of first two principal components of the AMMI analysis for heading date (Z55) of 151 spring two-rowed cultivars over 16 field trials.

### QTL analysis at single environments

GWA analyses of the four phenotypic traits at the single-trial level produced relatively weak associations. For grain yield, only two MTAs were detected above a Bonferroni threshold of P<0.05, while 20 MTAs were detected for heading date and none both for plant height and for thousand grain weight. Both MTAs of GY were detected from ITA10; the 20 MTAs for HD were from ESP16. Lowering the threshold to a more liberal p<0.0001, commonly used in GWA studies, still detected 125 MTAs for grain yield, 96 MTAs for heading date, 54 MTAs for plant height, and 10 MTAs for TGW (Table S3). Some regions with common QTLs across trials were suggested but, overall, the number and strength of associations found was low.

### QTL meta-analysis

The meta-analyses amplified the association signals, based on the relevance of *p*-values and the commonality in the direction of the effects sign across trials. The results were analysed using a stringent threshold based on permutations. Many SNPs had associations above that threshold. Some markers clearly indicated the same chromosome region. Associated markers were merged into QTLs, combining several criteria. All associated markers in a chromosome were subjected to cluster analysis, as in Looseley et al. (2020), suggesting groups likely belonging to the same QTL. The local LD decay (and chromosomal LD decay, if local LD was indeterminate), helped to delimit the QTLs on each chromosome. This process resulted in the detection of 23 QTLs for heading date, 29 for plant height, 11 for grain yield, and 27 for thousand grain weight (Table 3, Fig.2, Fig. S8). These numbers are rather high, because of the detection of QTLs having minor effects, which are usually not found in studies of smaller scale. The meta-analysis tends to identify MTAs that show the same sign across all trials. For this reason, it is not expected that it will capture qualitative QTL-by-environment interactions, if these exist. Therefore, most QTLs were rather consistent across trials (Table 3). However, some patterns of interaction were evident for a few QTLs, particularly for heading date. The differences in significance between the autumn- and spring-sown trials were large in some cases. The QTLs for HD5 (Fig. 3), HD6, and HD8 were much more significant in spring than in autumn trials. Conversely, HD21 was relatively more important in autumn-sown trials. There was a single QTL detected in the analyses of HD split by sowing time that was not detected in the global analysis. HDA1, on 1H, was significant only in the autumn-sown trials, indicating a marked QTL-by-environment interaction, possibly of qualitative nature, at this locus. The QTL-by-environment interaction was less evident for plant height (Table 3), as shown by PH14 (Fig. 3).

**Table 3.**
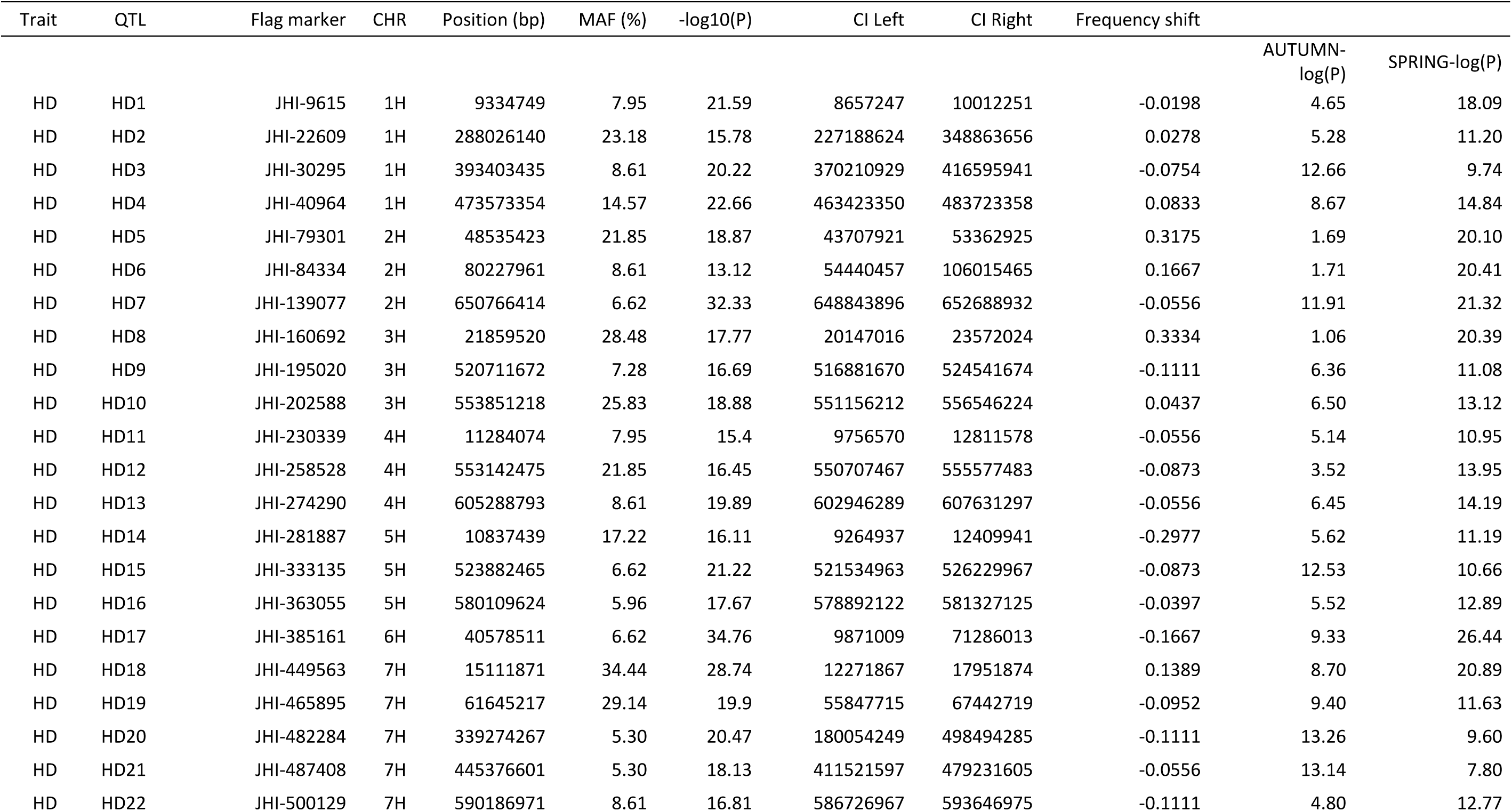
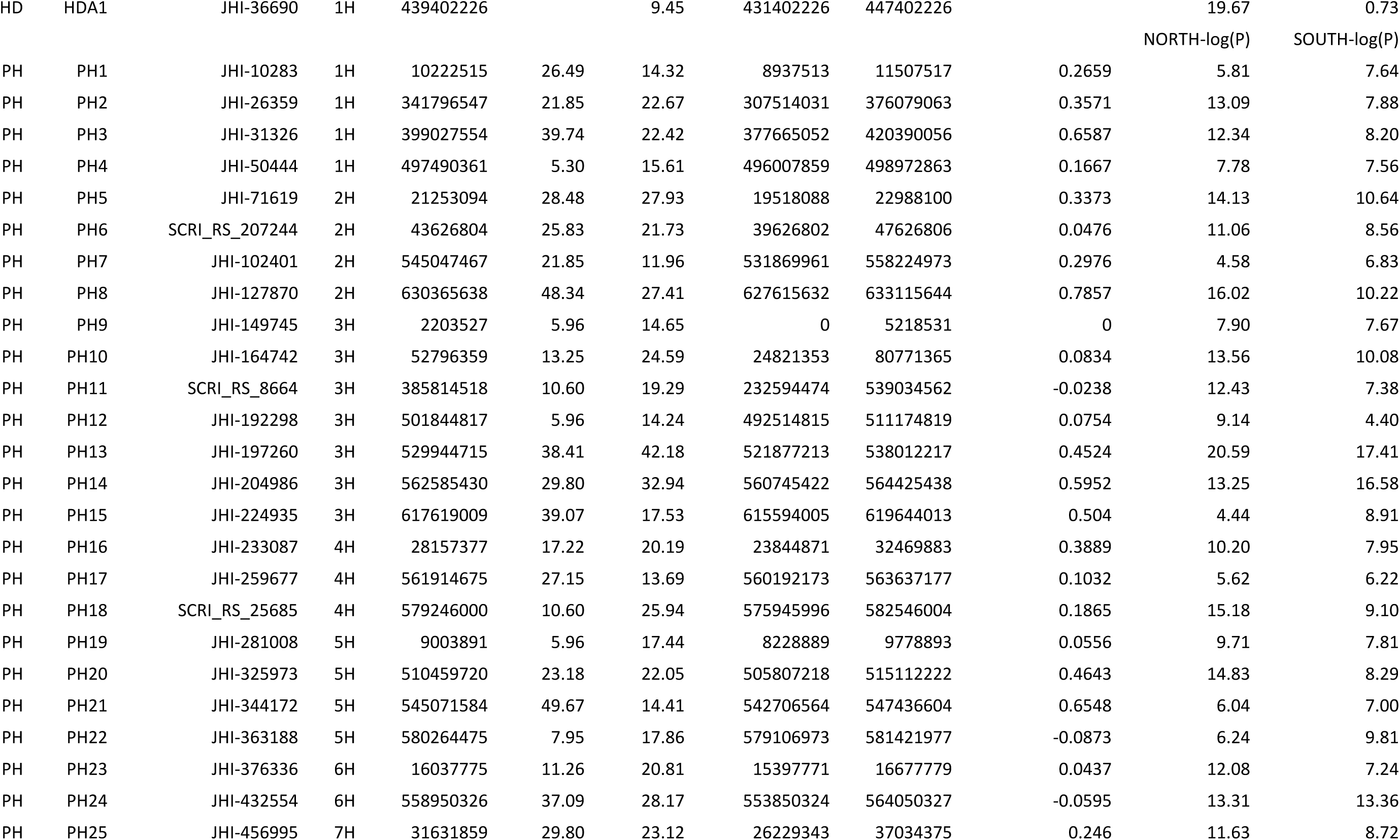
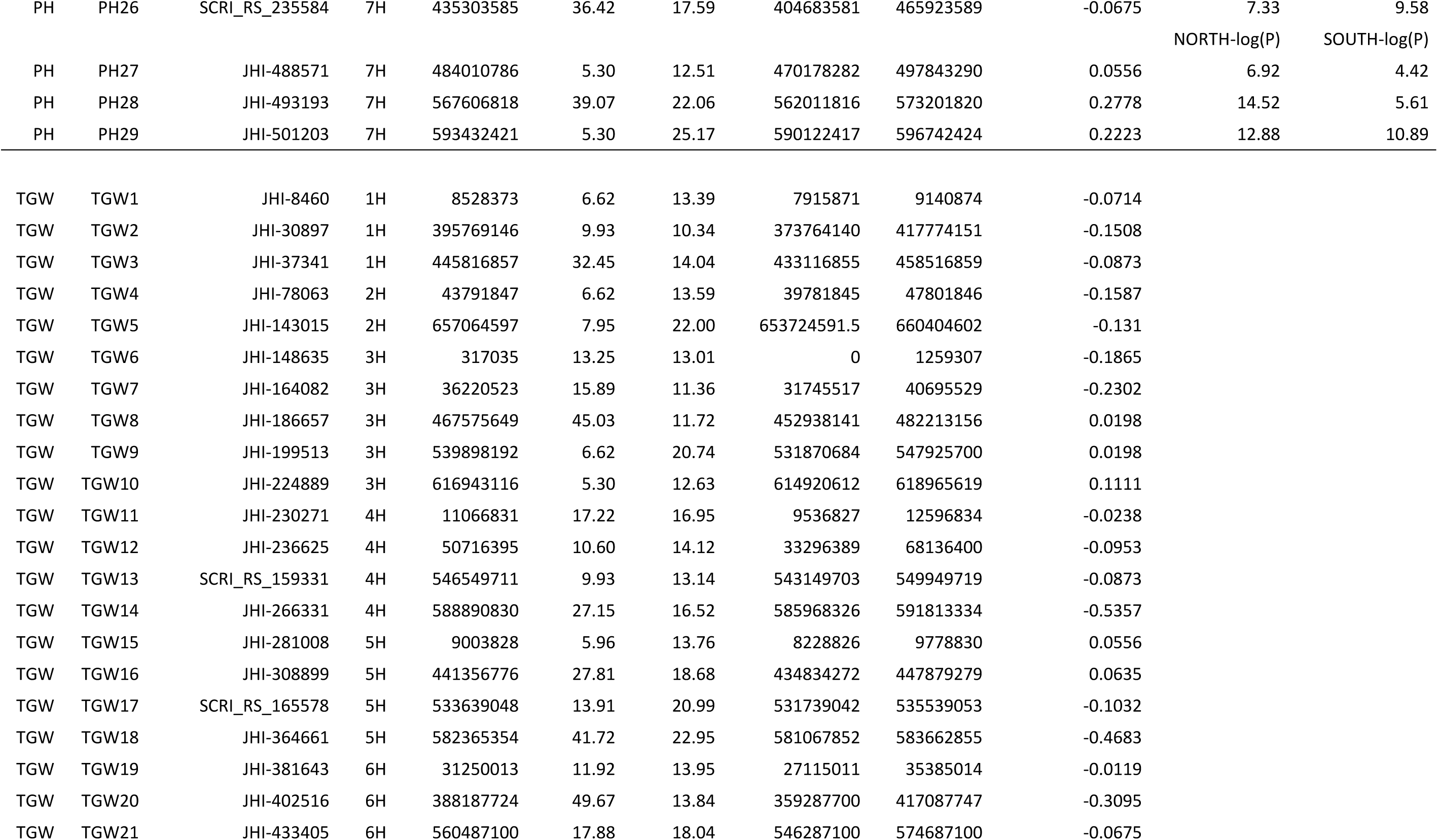
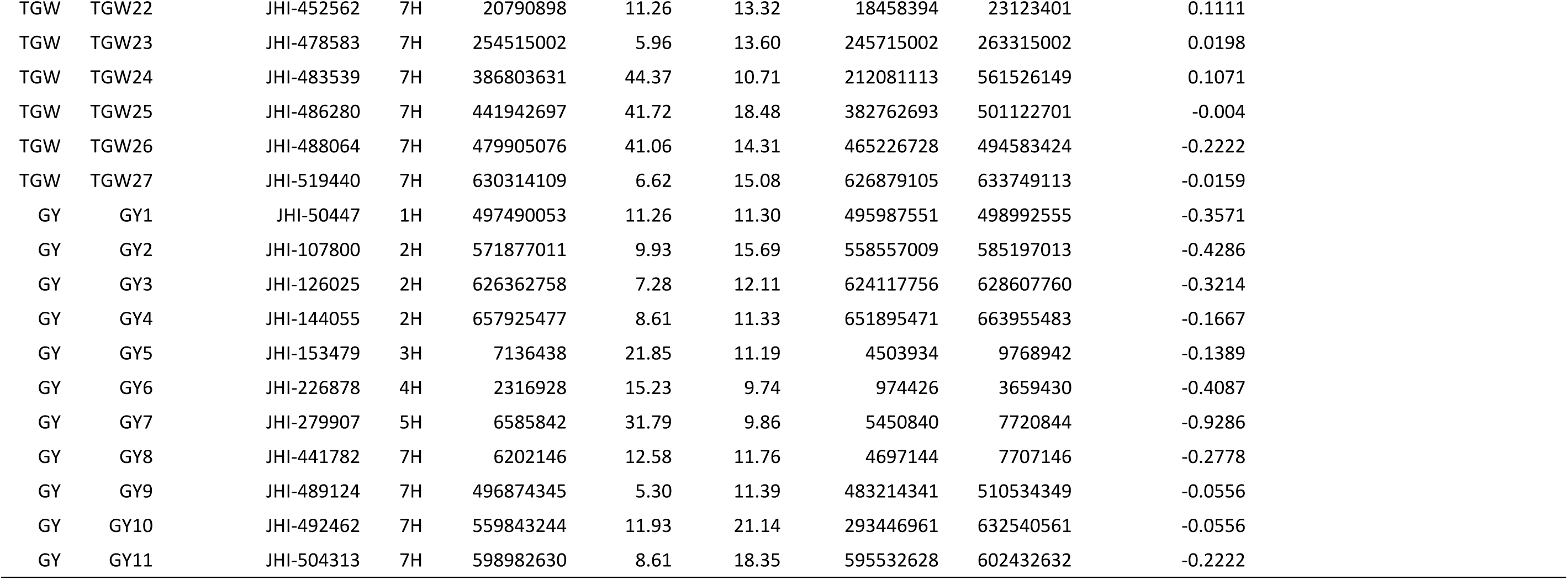
Codes, positions, and significance of QTLs detected in the meta-analyses for the four traits studied. The last two columns correspond to the meta-analyses carried out splitting the trials in two subsets, according to the main genotype-by-environment trends found for HD and PH.

| Trait | QTL | Flag marker | CHR | Position (bp) | MAF (%) | -log10(P) | CI Left | CI Right | Frequency shift | AUTUMN-log(P) | SPRING-log(P) |
| --- | --- | --- | --- | --- | --- | --- | --- | --- | --- | --- | --- |
| HD | HD1 | JHI-9615 | 1H | 9334749 | 7.95 | 21.59 | 8657247 | 10012251 | -0.0198 | 4.65 | 18.09 |
| HD | HD2 | JHI-22609 | 1H | 288026140 | 23.18 | 15.78 | 227188624 | 348863656 | 0.0278 | 5.28 | 11.20 |
| HD | HD3 | JHI-30295 | 1H | 393403435 | 8.61 | 20.22 | 370210929 | 416595941 | -0.0754 | 12.66 | 9.74 |
| HD | HD4 | JHI-40964 | 1H | 473573354 | 14.57 | 22.66 | 463423350 | 483723358 | 0.0833 | 8.67 | 14.84 |
| HD | HD5 | JHI-79301 | 2H | 48535423 | 21.85 | 18.87 | 43707921 | 53362925 | 0.3175 | 1.69 | 20.10 |
| HD | HD6 | JHI-84334 | 2H | 80227961 | 8.61 | 13.12 | 54440457 | 106015465 | 0.1667 | 1.71 | 20.41 |
| HD | HD7 | JHI-139077 | 2H | 650766414 | 6.62 | 32.33 | 648843896 | 652688932 | -0.0556 | 11.91 | 21.32 |
| HD | HD8 | JHI-160692 | 3H | 21859520 | 28.48 | 17.77 | 20147016 | 23572024 | 0.3334 | 1.06 | 20.39 |
| HD | HD9 | JHI-195020 | 3H | 520711672 | 7.28 | 16.69 | 516881670 | 524541674 | -0.1111 | 6.36 | 11.08 |
| HD | HD10 | JHI-202588 | 3H | 553851218 | 25.83 | 18.88 | 551156212 | 556546224 | 0.0437 | 6.50 | 13.12 |
| HD | HD11 | JHI-230339 | 4H | 11284074 | 7.95 | 15.4 | 9756570 | 12811578 | -0.0556 | 5.14 | 10.95 |
| HD | HD12 | JHI-258528 | 4H | 553142475 | 21.85 | 16.45 | 550707467 | 555577483 | -0.0873 | 3.52 | 13.95 |
| HD | HD13 | JHI-274290 | 4H | 605288793 | 8.61 | 19.89 | 602946289 | 607631297 | -0.0556 | 6.45 | 14.19 |
| HD | HD14 | JHI-281887 | 5H | 10837439 | 17.22 | 16.11 | 9264937 | 12409941 | -0.2977 | 5.62 | 11.19 |
| HD | HD15 | JHI-333135 | 5H | 523882465 | 6.62 | 21.22 | 521534963 | 526229967 | -0.0873 | 12.53 | 10.66 |
| HD | HD16 | JHI-363055 | 5H | 580109624 | 5.96 | 17.67 | 578892122 | 581327125 | -0.0397 | 5.52 | 12.89 |
| HD | HD17 | JHI-385161 | 6H | 40578511 | 6.62 | 34.76 | 9871009 | 71286013 | -0.1667 | 9.33 | 26.44 |
| HD | HD18 | JHI-449563 | 7H | 15111871 | 34.44 | 28.74 | 12271867 | 17951874 | 0.1389 | 8.70 | 20.89 |
| HD | HD19 | JHI-465895 | 7H | 61645217 | 29.14 | 19.9 | 55847715 | 67442719 | -0.0952 | 9.40 | 11.63 |
| HD | HD20 | JHI-482284 | 7H | 339274267 | 5.30 | 20.47 | 180054249 | 498494285 | -0.1111 | 13.26 | 9.60 |
| HD | HD21 | JHI-487408 | 7H | 445376601 | 5.30 | 18.13 | 411521597 | 479231605 | -0.0556 | 13.14 | 7.80 |
| HD | HD22 | JHI-500129 | 7H | 590186971 | 8.61 | 16.81 | 586726967 | 593646975 | -0.1111 | 4.80 | 12.77 |

| Trait | QTL | Flag marker | CHR | Position (bp) | MAF (%) | -log10(P) | CI Left | CI Right | Frequency shift |  |  |
| --- | --- | --- | --- | --- | --- | --- | --- | --- | --- | --- | --- |
| HD | HDA1 | JHI-36690 | 1H | 439402226 |  | 9.45 | 431402226 | 447402226 |  | 19.67 | 0.73 |
|  |  |  |  |  |  |  |  |  |  | NORTH-log(P) | SOUTH-log(P) |
| PH | PH1 | JHI-10283 | 1H | 10222515 | 26.49 | 14.32 | 8937513 | 11507517 | 0.2659 | 5.81 | 7.64 |
| PH | PH2 | JHI-26359 | 1H | 341796547 | 21.85 | 22.67 | 307514031 | 376079063 | 0.3571 | 13.09 | 7.88 |
| PH | PH3 | JHI-31326 | 1H | 399027554 | 39.74 | 22.42 | 377665052 | 420390056 | 0.6587 | 12.34 | 8.20 |
| PH | PH4 | JHI-50444 | 1H | 497490361 | 5.30 | 15.61 | 496007859 | 498972863 | 0.1667 | 7.78 | 7.56 |
| PH | PH5 | JHI-71619 | 2H | 21253094 | 28.48 | 27.93 | 19518088 | 22988100 | 0.3373 | 14.13 | 10.64 |
| PH | PH6 | SCRI_RS_207244 | 2H | 43626804 | 25.83 | 21.73 | 39626802 | 47626806 | 0.0476 | 11.06 | 8.56 |
| PH | PH7 | JHI-102401 | 2H | 545047467 | 21.85 | 11.96 | 531869961 | 558224973 | 0.2976 | 4.58 | 6.83 |
| PH | PH8 | JHI-127870 | 2H | 630365638 | 48.34 | 27.41 | 627615632 | 633115644 | 0.7857 | 16.02 | 10.22 |
| PH | PH9 | JHI-149745 | 3H | 2203527 | 5.96 | 14.65 | 0 | 5218531 | 0 | 7.90 | 7.67 |
| PH | PH10 | JHI-164742 | 3H | 52796359 | 13.25 | 24.59 | 24821353 | 80771365 | 0.0834 | 13.56 | 10.08 |
| PH | PH11 | SCRI_RS_8664 | 3H | 385814518 | 10.60 | 19.29 | 232594474 | 539034562 | -0.0238 | 12.43 | 7.38 |
| PH | PH12 | JHI-192298 | 3H | 501844817 | 5.96 | 14.24 | 492514815 | 511174819 | 0.0754 | 9.14 | 4.40 |
| PH | PH13 | JHI-197260 | 3H | 529944715 | 38.41 | 42.18 | 521877213 | 538012217 | 0.4524 | 20.59 | 17.41 |
| PH | PH14 | JHI-204986 | 3H | 562585430 | 29.80 | 32.94 | 560745422 | 564425438 | 0.5952 | 13.25 | 16.58 |
| PH | PH15 | JHI-224935 | 3H | 617619009 | 39.07 | 17.53 | 615594005 | 619644013 | 0.504 | 4.44 | 8.91 |
| PH | PH16 | JHI-233087 | 4H | 28157377 | 17.22 | 20.19 | 23844871 | 32469883 | 0.3889 | 10.20 | 7.95 |
| PH | PH17 | JHI-259677 | 4H | 561914675 | 27.15 | 13.69 | 560192173 | 563637177 | 0.1032 | 5.62 | 6.22 |
| PH | PH18 | SCRI_RS_25685 | 4H | 579246000 | 10.60 | 25.94 | 575945996 | 582546004 | 0.1865 | 15.18 | 9.10 |
| PH | PH19 | JHI-281008 | 5H | 9003891 | 5.96 | 17.44 | 8228889 | 9778893 | 0.0556 | 9.71 | 7.81 |
| PH | PH20 | JHI-325973 | 5H | 510459720 | 23.18 | 22.05 | 505807218 | 515112222 | 0.4643 | 14.83 | 8.29 |
| PH | PH21 | JHI-344172 | 5H | 545071584 | 49.67 | 14.41 | 542706564 | 547436604 | 0.6548 | 6.04 | 7.00 |
| PH | PH22 | JHI-363188 | 5H | 580264475 | 7.95 | 17.86 | 579106973 | 581421977 | -0.0873 | 6.24 | 9.81 |
| PH | PH23 | JHI-376336 | 6H | 16037775 | 11.26 | 20.81 | 15397771 | 16677779 | 0.0437 | 12.08 | 7.24 |
| PH | PH24 | JHI-432554 | 6H | 558950326 | 37.09 | 28.17 | 553850324 | 564050327 | -0.0595 | 13.31 | 13.36 |
| PH | PH25 | JHI-456995 | 7H | 31631859 | 29.80 | 23.12 | 26229343 | 37034375 | 0.246 | 11.63 | 8.72 |
| PH | PH26 | SCRI_RS_235584 | 7H | 435303585 | 36.42 | 17.59 | 404683581 | 465923589 | -0.0675 | 7.33 | 9.58 |
|  |  |  |  |  |  |  |  |  |  | NORTH-log(P) | SOUTH-log(P) |
| PH | PH27 | JHI-488571 | 7H | 484010786 | 5.30 | 12.51 | 470178282 | 497843290 | 0.0556 | 6.92 | 4.42 |
| PH | PH28 | JHI-493193 | 7H | 567606818 | 39.07 | 22.06 | 562011816 | 573201820 | 0.2778 | 14.52 | 5.61 |
| PH | PH29 | JHI-501203 | 7H | 593432421 | 5.30 | 25.17 | 590122417 | 596742424 | 0.2223 | 12.88 | 10.89 |
| TGW | TGW1 | JHI-8460 | 1H | 8528373 | 6.62 | 13.39 | 7915871 | 9140874 | -0.0714 |  |  |
| TGW | TGW2 | JHI-30897 | 1H | 395769146 | 9.93 | 10.34 | 373764140 | 417774151 | -0.1508 |  |  |
| TGW | TGW3 | JHI-37341 | 1H | 445816857 | 32.45 | 14.04 | 433116855 | 458516859 | -0.0873 |  |  |
| TGW | TGW4 | JHI-78063 | 2H | 43791847 | 6.62 | 13.59 | 39781845 | 47801846 | -0.1587 |  |  |
| TGW | TGW5 | JHI-143015 | 2H | 657064597 | 7.95 | 22.00 | 653724591.5 | 660404602 | -0.131 |  |  |
| TGW | TGW6 | JHI-148635 | 3H | 317035 | 13.25 | 13.01 | 0 | 1259307 | -0.1865 |  |  |
| TGW | TGW7 | JHI-164082 | 3H | 36220523 | 15.89 | 11.36 | 31745517 | 40695529 | -0.2302 |  |  |
| TGW | TGW8 | JHI-186657 | 3H | 467575649 | 45.03 | 11.72 | 452938141 | 482213156 | 0.0198 |  |  |
| TGW | TGW9 | JHI-199513 | 3H | 539898192 | 6.62 | 20.74 | 531870684 | 547925700 | 0.0198 |  |  |
| TGW | TGW10 | JHI-224889 | 3H | 616943116 | 5.30 | 12.63 | 614920612 | 618965619 | 0.1111 |  |  |
| TGW | TGW11 | JHI-230271 | 4H | 11066831 | 17.22 | 16.95 | 9536827 | 12596834 | -0.0238 |  |  |
| TGW | TGW12 | JHI-236625 | 4H | 50716395 | 10.60 | 14.12 | 33296389 | 68136400 | -0.0953 |  |  |
| TGW | TGW13 | SCRI_RS_159331 | 4H | 546549711 | 9.93 | 13.14 | 543149703 | 549949719 | -0.0873 |  |  |
| TGW | TGW14 | JHI-266331 | 4H | 588890830 | 27.15 | 16.52 | 585968326 | 591813334 | -0.5357 |  |  |
| TGW | TGW15 | JHI-281008 | 5H | 9003828 | 5.96 | 13.76 | 8228826 | 9778830 | 0.0556 |  |  |
| TGW | TGW16 | JHI-308899 | 5H | 441356776 | 27.81 | 18.68 | 434834272 | 447879279 | 0.0635 |  |  |
| TGW | TGW17 | SCRI_RS_165578 | 5H | 533639048 | 13.91 | 20.99 | 531739042 | 535539053 | -0.1032 |  |  |
| TGW | TGW18 | JHI-364661 | 5H | 582365354 | 41.72 | 22.95 | 581067852 | 583662855 | -0.4683 |  |  |
| TGW | TGW19 | JHI-381643 | 6H | 31250013 | 11.92 | 13.95 | 27115011 | 35385014 | -0.0119 |  |  |
| TGW | TGW20 | JHI-402516 | 6H | 388187724 | 49.67 | 13.84 | 359287700 | 417087747 | -0.3095 |  |  |
| TGW | TGW21 | JHI-433405 | 6H | 560487100 | 17.88 | 18.04 | 546287100 | 574687100 | -0.0675 |  |  |

| Trait | QTL | Flag marker | CHR | Position (bp) | MAF (%) | -log10(P) | CI Left | CI Right | Frequency shift |
| --- | --- | --- | --- | --- | --- | --- | --- | --- | --- |
| TGW | TGW22 | JHI-452562 | 7H | 20790898 | 11.26 | 13.32 | 18458394 | 23123401 | 0.1111 |
| TGW | TGW23 | JHI-478583 | 7H | 254515002 | 5.96 | 13.60 | 245715002 | 263315002 | 0.0198 |
| TGW | TGW24 | JHI-483539 | 7H | 386803631 | 44.37 | 10.71 | 212081113 | 561526149 | 0.1071 |
| TGW | TGW25 | JHI-486280 | 7H | 441942697 | 41.72 | 18.48 | 382762693 | 501122701 | -0.004 |
| TGW | TGW26 | JHI-488064 | 7H | 479905076 | 41.06 | 14.31 | 465226728 | 494583424 | -0.2222 |
| TGW | TGW27 | JHI-519440 | 7H | 630314109 | 6.62 | 15.08 | 626879105 | 633749113 | -0.0159 |
| GY | GY1 | JHI-50447 | 1H | 497490053 | 11.26 | 11.30 | 495987551 | 498992555 | -0.3571 |
| GY | GY2 | JHI-107800 | 2H | 571877011 | 9.93 | 15.69 | 558557009 | 585197013 | -0.4286 |
| GY | GY3 | JHI-126025 | 2H | 626362758 | 7.28 | 12.11 | 624117756 | 628607760 | -0.3214 |
| GY | GY4 | JHI-144055 | 2H | 657925477 | 8.61 | 11.33 | 651895471 | 663955483 | -0.1667 |
| GY | GY5 | JHI-153479 | 3H | 7136438 | 21.85 | 11.19 | 4503934 | 9768942 | -0.1389 |
| GY | GY6 | JHI-226878 | 4H | 2316928 | 15.23 | 9.74 | 974426 | 3659430 | -0.4087 |
| GY | GY7 | JHI-279907 | 5H | 6585842 | 31.79 | 9.86 | 5450840 | 7720844 | -0.9286 |
| GY | GY8 | JHI-441782 | 7H | 6202146 | 12.58 | 11.76 | 4697144 | 7707146 | -0.2778 |
| GY | GY9 | JHI-489124 | 7H | 496874345 | 5.30 | 11.39 | 483214341 | 510534349 | -0.0556 |
| GY | GY10 | JHI-492462 | 7H | 559843244 | 11.93 | 21.14 | 293446961 | 632540561 | -0.0556 |
| GY | GY11 | JHI-504313 | 7H | 598982630 | 8.61 | 18.35 | 595532628 | 602432632 | -0.2222 |

**Figure 2.**
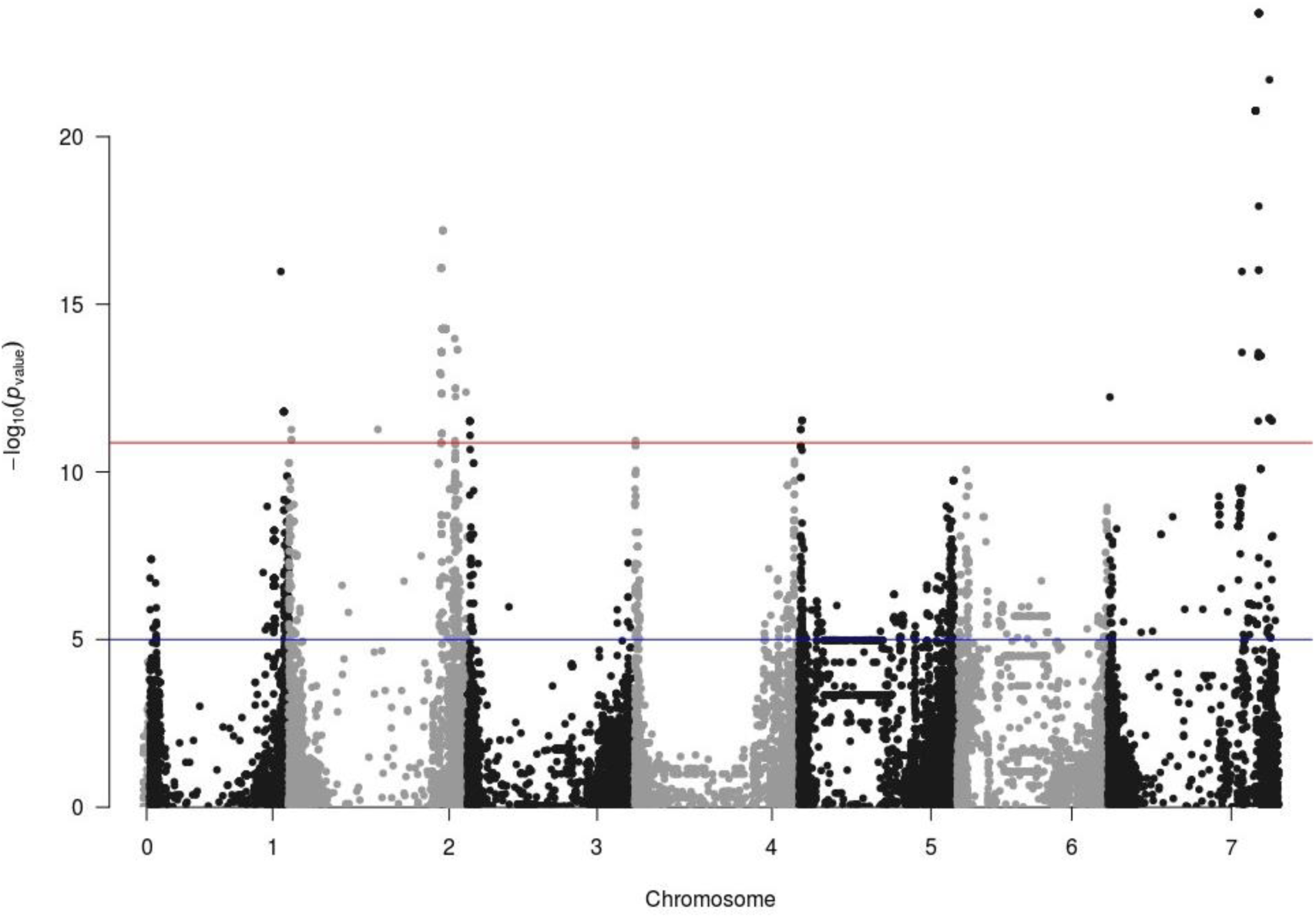
Manhattan plot of the meta-analysis of 14 trials for grain yield (GY). In blue, threshold commonly used in meta-analyses in the literature. In red, threshold calculated in this study, corresponding to the minimum P-value resulting from 1000 permutations.

**Figure 3.**
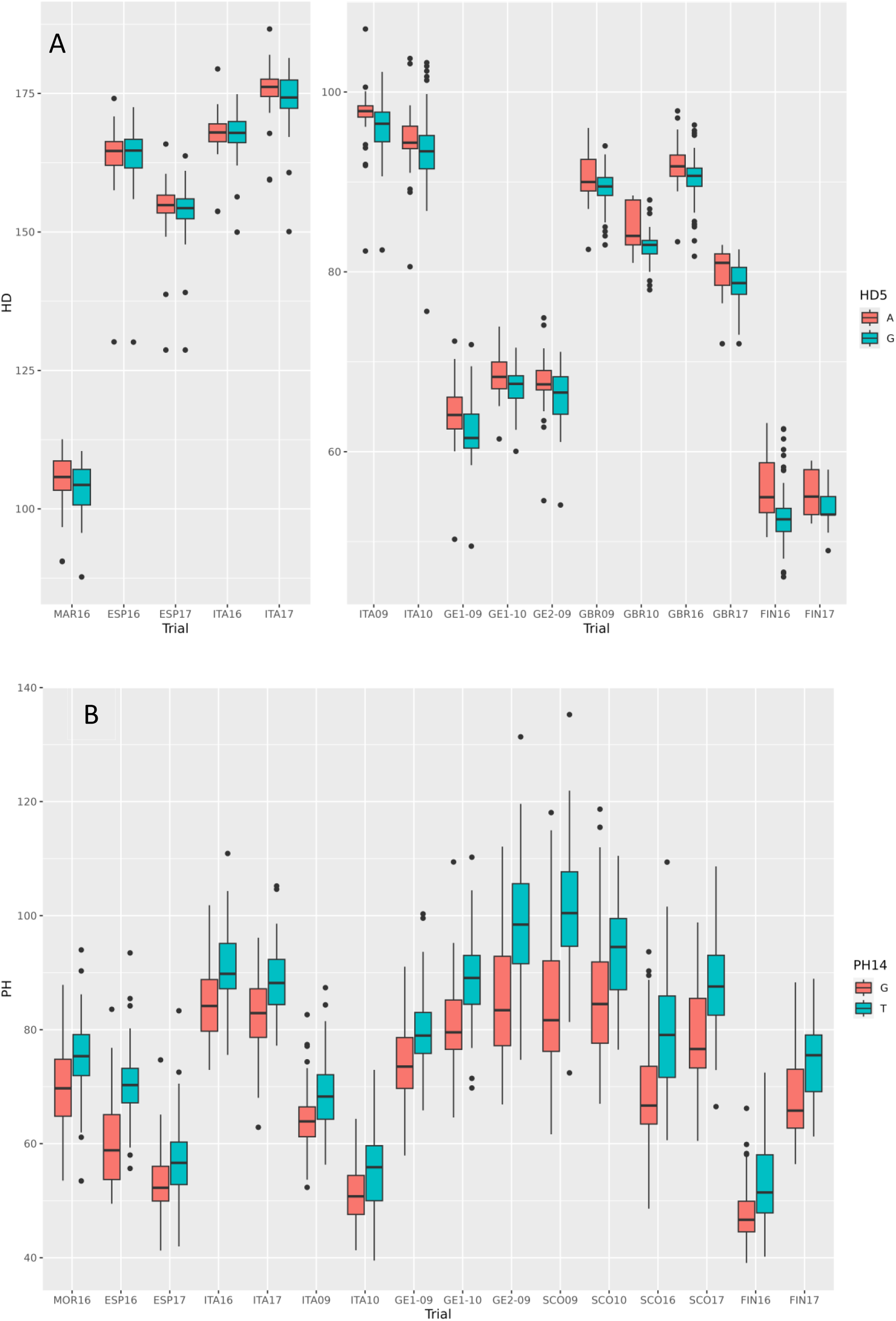
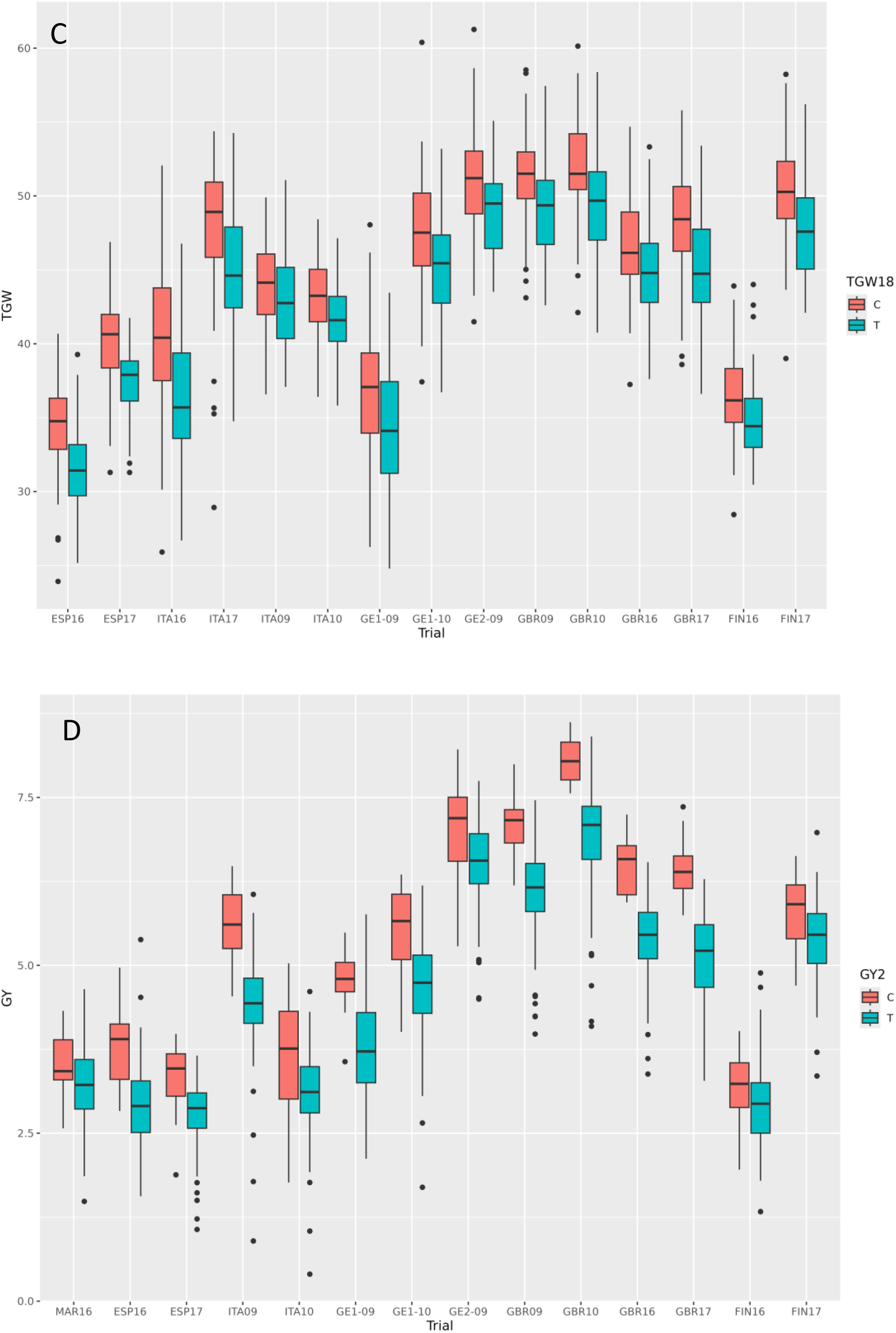
Allelic boxplots for four QTL (one for each trait), across environments. A) QTL HD5, days from sowing to heading, divided into autumn (left panel) and winter-spring sowings (right panel); B) QTL PH14, plant height; C) QTL TGW18, thousand grain weight; D) QTL GY2, grain yield.

Multilocus models derived from sequential MANOVA analyses were used to identify the best subset of significant QTLs that jointly explained each trait. All QTLs included in the models were significant for P<0.05 or less and had partial Eta squared (ƞp²) above 0.14. This value is commonly used as threshold to declare the influence of independent factors on the dependent variables. These joint models explained around 16% of phenotypic variance for HD, TGW, and GY, and 33% for PH (Table 4). The QTLs retained in the models had a rather small effect, explaining from 1.08 up to 6.07 percent of the variance of the traits: from 0.14 to 0.33 t ha^-1^ (averaged across all environments) for GY, from 0.7 to 1.9 days for HD, 1.4 to 2.7 cm for PH, and 0.6 to 1.3 g for TGW. The QTLs explaining more than 5% of the variance of the trait were PH28, on 7H (2.5 cm), and GY2, on 1H (5.08%, 0.28 t ha^-1^).

**Table 4.** Summary of QTLs kept in the multivariate multilocus analysis for the four studied traits.

| Trait | Chr | QTL | Best model | Partial Eta2 | Percentage of explained variance |
| --- | --- | --- | --- | --- | --- |
| HD | 1H | HD1 | * | 0.20 | 1.39 |
| HD | 2H | HD5 | *** | 0.37 | 2.89 |
| HD | 3H | HD8 | *** | 0.30 | 2.23 |
| HD | 5H | HD16 | *** | 0.22 | 1.58 |
| HD | 6H | HD17 | *** | 0.26 | 1.84 |
| HD | 7H | HD18 | *** | 0.27 | 1.95 |
| HD | 7H | HD21 | *** | 0.40 | 3.17 |
| HD | 7H | HD22 | * | 0.21 | 1.47 |
| Total HD: 16.52 |  |  |  |  |  |
| PH | 1H | PH1 | *** | 0.40 | 3.17 |
| PH | 2H | PH6 | *** | 0.30 | 2.18 |
| PH | 2H | PH8 | *** | 0.30 | 2.23 |
| PH | 3H | PH11 | ** | 0.26 | 1.89 |
| PH | 3H | PH13 | ** | 0.23 | 1.58 |
| PH | 3H | PH14 | *** | 0.43 | 3.45 |
| PH | 4H | PH16 | ** | 0.23 | 1.65 |
| PH | 4H | PH17 | * | 0.21 | 1.48 |
| PH | 5H | PH20 | *** | 0.54 | 4.72 |
| PH | 5H | PH21 | ** | 0.23 | 1.66 |
| PH | 6H | PH23 | * | 0.20 | 1.42 |
| PH | 7H | PH26 | * | 0.21 | 1.50 |
| PH | 7H | PH28 | *** | 0.63 | 6.07 |
| Total PH: 32.99 |  |  |  |  |  |
| TGW | 1H | TGW3 | * | 0.18 | 1.22 |
| TGW | 2H | TGW5 | ** | 0.23 | 1.63 |
| TGW | 4H | TGW14 | *** | 0.39 | 3.02 |
| TGW | 5H | TGW15 | *** | 0.32 | 2.40 |
| TGW | 5H | TGW16 | * | 0.20 | 1.36 |
| TGW | 5H | TGW18 | *** | 0.26 | 1.85 |
| TGW | 6H | TGW20 | * | 0.18 | 1.23 |
| TGW | 7H | TGW26 | *** | 0.32 | 2.41 |
| TGW | 7H | TGW27 | ** | 0.21 | 1.50 |
| Total TGW: 16.61 |  |  |  |  |  |
| GY | 2H | GY2 | *** | 0.57 | 5.08 |
| GY | 2H | GY4 | ** | 0.20 | 1.41 |
| GY | 4H | GY6 | . | 0.16 | 1.08 |
| GY | 5H | GY7 | *** | 0.30 | 2.23 |
| GY | 7H | GY9 | *** | 0.26 | 1.86 |
| GY | 7H | GY10 | * | 0.18 | 1.22 |
| GY | 7H | GY11 | *** | 0.39 | 3.00 |
| Total GY: 15.88 |  |  |  |  |  |

The intervals for confidence regions of the QTLs were wide, with a mean size of 25.95 Mb. Regions ranged from 1.23 Mb to 349.45 Mb, illustrating just how unevenly distributed is LD in spring barley germplasm. Moreover, two regions of nearly 350 Mb were identified on chromosome 7H, most likely related to an inversion of 141 Mb (Jayakodi et al., 2020) that already has been identified in some cultivars included in our association panel. There was overlap between the confidence intervals for QTLs of different traits, as indicated in Table S4. The region of TGW15-PH19 could represent the same QTL with pleiotropic effects. Some grain yield QTLs overlap with plant height (PH4), and thousand grain weight (TGW5, TGW25, TGW26, and TGW27). Other QTLs were linked to some extent, which may have implications for their management in breeding.

### Temporal variation in traits and QTL allelic frequencies

The classification of cultivars into four classes, according to the year of release, revealed salient trait and QTL trends resulting from breeding efforts and preferences. Plant height and thousand-grain weight presented marked trends towards reduced (PH) or increased (TGW) values through the years, in accordance with the expected outcome from breeding programs (Fig. 4) and these traits’ heritability (Table 1). The trends were highly consistent across environments (Fig. S4). For grain yield, the increase over time was less marked, except in Scotland. In general, the largest yield improvement was observed for the most modern cultivars compared to earlier ones. There were no marked historic trends regarding heading date. The only remarkable feature was that the most modern class of cultivars was always the earliest in all spring-grown trials, whereas this difference was not observed in the autumn-sown trials, hinting at the occurrence of a possible genotype-by-environment interaction.

**Figure 4.**
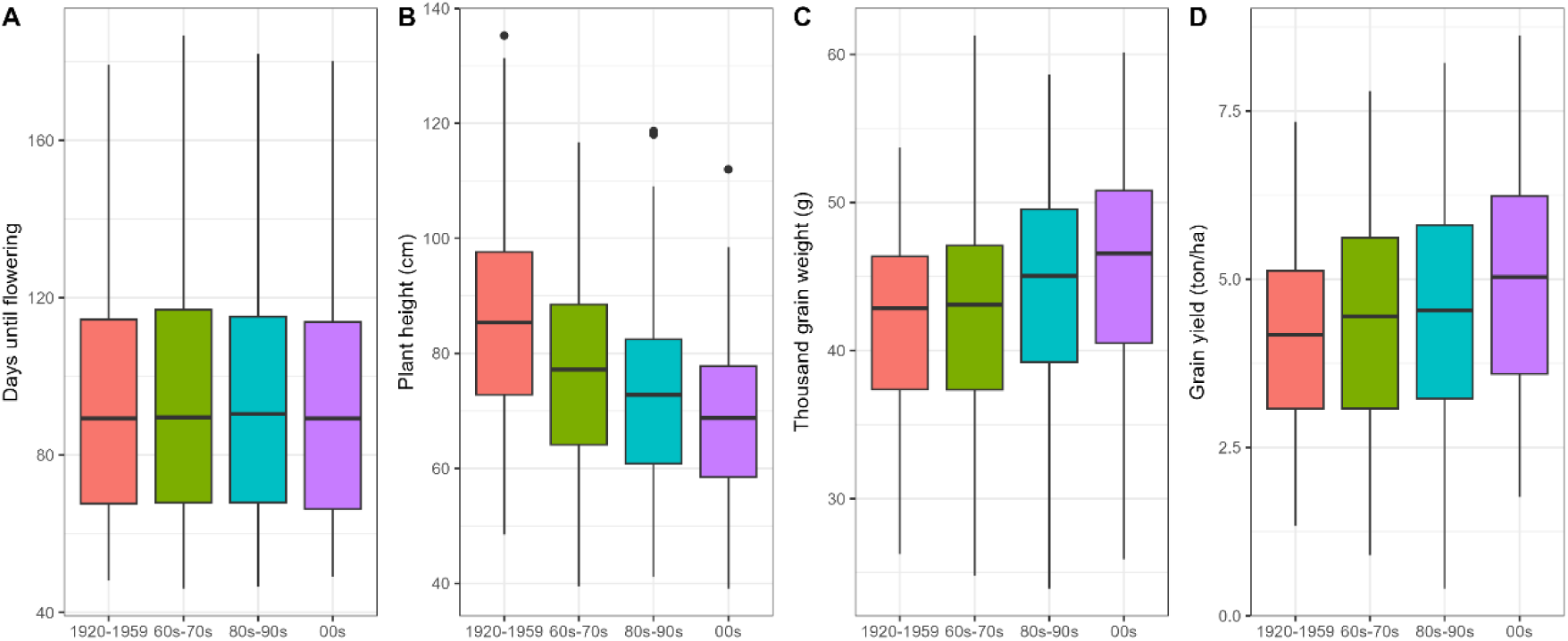
Phenotype boxplots of all trials divided by groups of release years. A) Days from sowing to heading, B) plant height, C) thousand grain weight and D) grain yield.

Regarding allelic frequencies for QTLs over time, some QTLs followed the general trend for their trait, while others showed no apparent response. For PH, the height-increasing alleles of eight QTLs were already at low frequencies in the old cultivars; these remained low thereafter. Five QTLs (PH10, PH11, PH17, PH24, PH26) seemed not to be affected by selection, keeping high to very high frequencies of height-increasing alleles, whereas 11 QTLs exhibited marked increases in the frequency of height-reducing alleles (coloured lines, Fig. 5). For TGW (Fig. S9), the situation was similar, but with fewer QTLs apparently affected by selection. Five QTLs (TGW2, TGW5, TGW6, TGW11, TGW12) showed high frequencies of grain-weight-increasing alleles already in the oldest cultivar class, increasing slightly to almost fixation in the most modern class. The frequencies of another five QTLs (TGW7, TGW14, TGW18, TGW20, TGW26) for higher grain weight approximately doubled over time, i.e., were apparently favoured by selection. For another 17 QTLs, frequencies of favourable alleles varied between medium to low, with little or no response to selection. For GY, the frequency shifts of several QTLs occurred only in the most recent groups of cultivars. An exception was GY7, which was the QTL having the largest frequency change over time, considering all four analysed traits. For HD (Fig. S9), there were three QTLs that were apparently affected by selection (HD5, HD6 and HD8). These QTLs suffered some selection towards earliness. It is remarkable that these three QTLs were detected in the spring-sown trials, yet not in the autumn-sown ones (Table 3), showing the largest difference in significance between the two sets of trials. They may be responsible for the increased earliness shown by the most modern class of cultivars, which is seen only under spring-sown conditions.

**Figure 5.**
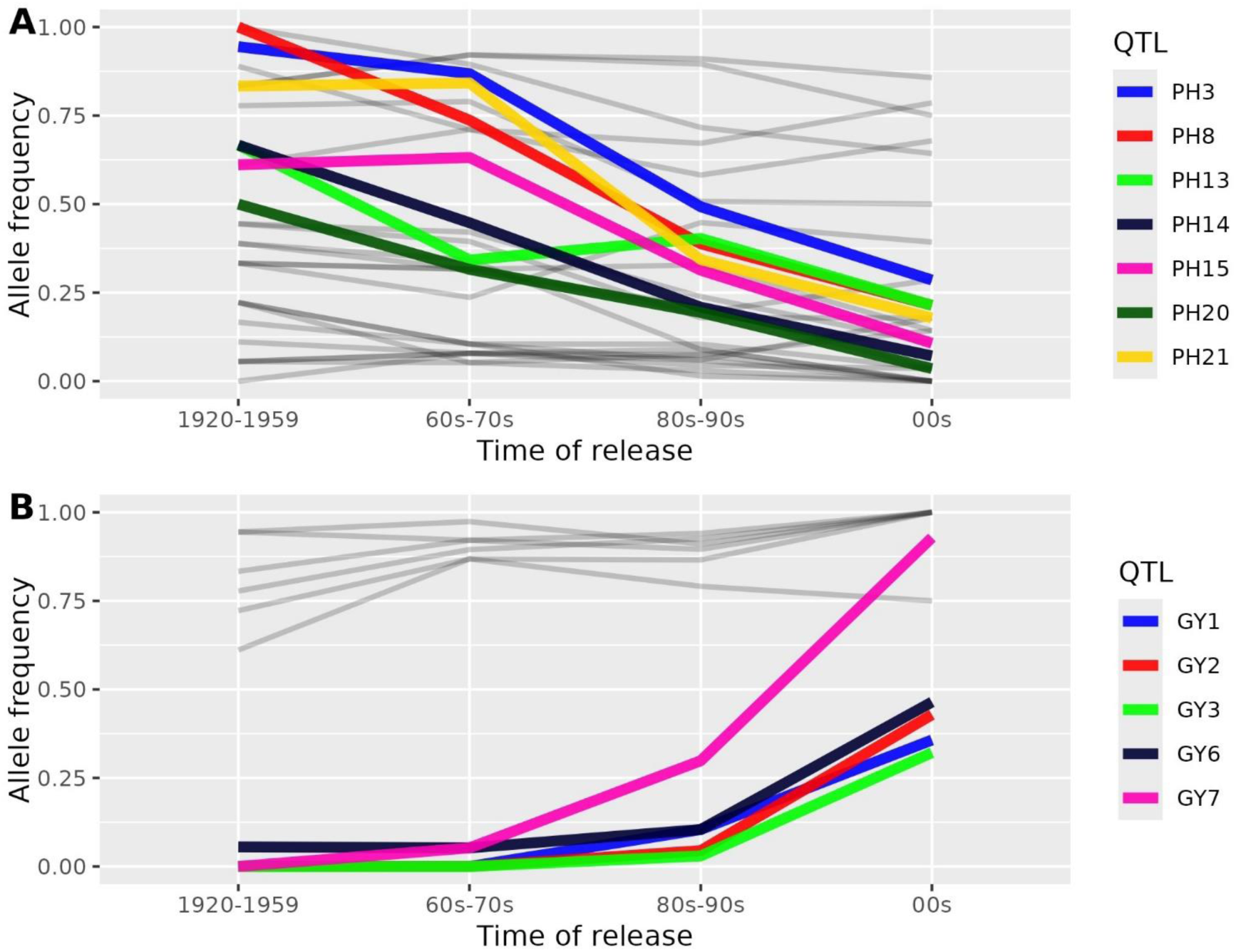
Changes in allele frequencies of the QTLs identified in the meta-analyses for plant height (A) and grain yield (B) across time of release of the cultivars.

At the genome level, some regions seemed preferentially targeted by selection. This was examined by looking at the difference in frequencies between the oldest and newest cultivar classes, calculated for the SNPs divided in 250 rolling windows (bins) per chromosome for all four groups of year of release (Fig. S10). Focusing on the 1% of bins with the largest frequency changes, the most visible selection footprint was in the pericentromeric area of chromosome 5H (65-310 Mb), coincident with the haplotype already detected by Wonneberger et al. (2023). Another narrow region in 5HL (around 543 Mb) showed similar allele frequency changes. Finally, a region in 4HL (510-530 Mb) also showed two close, narrow peaks of very large frequency shifts over time. For all traits, we have identified some QTLs apparently untouched by selection and some at low frequencies in the set studied. These could be aimed at by current breeders, provided they have not already been targeted by breeding in recent years and that they do not convey negative pleiotropic effects.

### Exome capture enrichment and candidate genes

The inclusion of exome capture (EC) data for the analysis of the QTLs detected provided 119,811 more markers. In 25 cases (31.65%), exome capture markers had equal or higher association than for the 50k SNPs (Table S5); these were designated as the new flag SNPs. For each QTL, a search of candidate genes for the flag markers was carried out within the confidence interval regions. In some cases, flag markers were present inside genes having annotations relevant for the traits considered. The EC markers provided higher resolution than did SNPs from the 50k set, for example, in the region of flowering time QTL HD17 on chromosome 6H. That confidence interval region harbours two flowering-related genes, *HvCMF3* (Cockram et al., 2012) and *HvZTLb* (Russell et al., 2016). However, the EC marker with the largest association is located inside gene model HORVU.MOREX.r3.6HG0557980, annotated as the nuclear pore complex protein Nup 160.

Several plant height QTLs presented EC markers with higher associations than those of markers in the 50k SNP set. PH1 showed highly associated EC markers in two different genes. While the most strongly associated marker was within a low-confidence gene, the second group of highly associated markers pointed to a nitrate transporter (HORVU.MOREX.r3.1HG0005090), which is an ortholog of rice *OsNRT1.4* (Bucher et al., 2014). PH24, which presents one of the highest associations, was within gene HORVU.MOREX.r3.6HG0632820, annotated as a WRKY transcription factor-like protein. The *sdw1* gene is a good candidate for QTL PH14. Mutations in this gene, a gibberellin 20-oxidase gene (*HvGA20ox2*), are one of the most common causes of semi-dwarfism in barley. It is a multiallelic locus, with a few alleles commonly employed in barley breeding to reduce plant height (Xu et al., 2017).

Half of the QTLs for thousand-grain weight contained interesting candidate genes, indicated by a higher association of an EC marker. For TGW18, included in the multivariate multilocus model, an ortholog of rice gene *OsBRXL4* was identified (HORVU.MOREX.r3.5HG0535760). The EC flag maker gene for TGW27 is HORVU.MOREX.r3.7HG0752360, which is a lysine-specific demethylase. Interestingly, this is the same annotation of the *Vrs3* gene (Bull et al., 2017; van Esse et al., 2017) and is expressed in developing inflorescences and grains. Its homologous gene in *A. thaliana* is activated under dehydration stress (Huang et al., 2019). For grain yield, enrichment with EC markers identified a gene coding an ɑ-glucosidase enzyme in QTL GY5. This gene, *HvAGL2* (HORVU.MOREX.r3.3HG0221900) was previously confirmed to be involved in starch metabolism in developing grains in barley (Andriotis et al., 2016). The GY2 confidence interval includes the *HvHOX1* (*Vrs1*) gene, involved in spike-row determination (Komatsuda et al., 2007). We used a specifically-designed KASP marker (Table S1) to genotype the panel for the *deficiens* allele *Vrs1.t.* None of the cultivars with the unfavourable allele at GY2 carried *Vrs1.t*. Out of the 15 cultivars carrying the favourable allele at QTL GY2, 14 were available to phenotype and genotype. Thirteen of them (all but Forum) carried the *deficiens* allele. An evaluation of the spikes of the 14 revealed differences in the size of the lateral spikelets. Eleven cultivars were phenotypically *deficiens*, without lateral spikelets. Forum was clearly not *deficiens*, genetically and phenotypically. Although Felicitas and Tocada are genetically *deficiens*, both showed very small laterals spikelets (Fig. S11). These results support *Vrs1* as a candidate gene for GY2.

## DISCUSSION

The sensitivity of the meta-analysis varied among traits. Using the rather liberal threshold (widely found in the literature) of –log10(*p*-value) = 4 at single-trial level, a total of 96 MTAs were detected for HD, in approximately 16 regions, with no QTL detected in more than three trials, while the meta-GWA allowed identification of 22 QTLs in high-confidence regions for HD (reduced to eight in the multilocus analysis). For PH and TGW, only three and six regions, respectively, were detected in single trials (with a maximum coincidence of five trials), much less than those identified by the meta-analysis. For grain yield, there were more regions detected in single trials, altogether 125 MTAs in 20 regions, compared to the 11 QTLs found in the meta-analysis. However, with a threshold of five, the number of MTAs dropped to 48 (in two QTL regions), zero, six (in one QTL), and 34 (in nine QTL regions), for HD, TGW, PH, and GY, respectively.

Overall, the meta-analysis increased the power of QTL detection; the confidence on the main-effect QTLs, particularly those in the multilocus analyses, is high. The more trials involved, the greater power of detection achieved. We were able to detect QTL effects as low as just above 1% of the traits’ variances. The stringent permutation test which was used protects against false positives. A strict LD control to determine the confidence intervals, and the use of multilocus models, restricted the number of QTLs detected even further, but we still found a high number of QTLs. Besides the increased power of the meta-analysis, breaking down the analysis by the main G x E component detected for HD and PH revealed QTLs and candidate genes specific for certain conditions, such as sowing date (autumn/spring) or latitude. The unique behaviour of cultivars Mona and Saana is likely caused by their carrying the same mutant allele in a major flowering time gene, *HvELF3* (Faure et al., 2012, Göransson et al., 2021). This causes them to bypass the photoperiod sensitivity mechanism, leading to early flowering irrespective of the daylength. This fact makes them relatively early in Southern environments, in which most of the growing period occurs under short photoperiod.

The QTL regions found in this study were compared with those of recent studies involving large populations and SNP genotyping, therefore allowing direct comparison. First, the co-localisation of plant height QTLs with those reported by Tondelli et al. (2013) was rather high. This is not surprising, as our genotype set is a subsample of theirs (they used a 216 spring two-rowed panel), and some trials were also shared (those from 2009 and 2010). We found 29 PH QTLs, compared with 17 found in Tondelli et al. (2013). Out of those 17 PH QTLs, 11 were inside the confidence intervals of our QTLs (PH1, PH2, PH4, PH6, PH11, PH13, PH14, or very close to them PH15, PH20, PH25, PH29). Plant height QTL derived from part of our trials (2016 and 2017) were reported by Bretani et al. (2022). They studied a larger diversity panel, comprising 165 two-rowed (including our entire panel), and 96 six-rowed barleys. They found 48 PH QTL, 26 of them either in the two-rowed or in the whole panel. Twelve of those 26 were inside the confidence intervals of nine of our QTLs (PH2, PH3, PH4, PH5, PH8, PH13, PH14, PH20, PH28). None of our QTLs coincided with their 22 six-rowed specific QTLs.

The same association panel as Tondelli et al. (2013) was studied by Xu et al. (2018). In this case, only two QTLs, one for grain yield (GY6) and one for thousand grain weight (TGW18), are shared with our results. This low number of matches could be explained by different methodologies for QTL detection. Multilocation QTL analysis was run using Genstat software, which allows finding QTL in interactions with the environment. As mentioned above, the meta-analysis focuses on QTLs with low interaction. Out of the 23 QTLs detected for GY and TGW in the work by Xu et al. (2018), only eight were detected as the main factor QTL (i.e., not interacting with the environment) and one of them coincided with a TGW found in our study.

Bustos-Korts et al. (2019) studied a highly diverse panel of 371 genotypes including landraces and cultivars, two- and six-rowed, winter and spring barley. They scored plant height, thousand grain weight, and flowering time (Z55) in common with our study. Two QTLs are shared for flowering time (HD12 and HD13), one of them in the region of *HvVRN2*. For plant height, two QTLs have matching confidence intervals (PH8 and PH14); *sdw1* is within one of them. For TGW, there is no QTL in common, although they found a QTL for this trait close to the *Vrs1* gene, which is the same location as our QTL GY2.

Recently, a large association panel (n = 363) of European two-rowed spring barley was analysed for flowering time, plant height, and thousand grain weight (Bernád et al., 2024). One QTL for flowering time on chromosome 6H had overlapping confidence intervals with HD17. Meanwhile, one MTA for plant height is within PH14, where *sdw1* is located, while another one on chromosome 7H is close to PH25, where their LD block should overlap. For thousand grain weight, one of their four QTLs has also been identified in this study (TGW9, chromosome 3H).

We found evidence of surprisingly highly colocalizing QTLs in a recent study addressing two- and six-rowed spring barley germplasm, which was less related to our panel than that of other previous studies. Two articles published using a diverse multi-parent barley population found new QTLs with minor to moderate phenotypic effects (Shrestha et al., 2022; Cosenza et al., 2024). Several of our HD QTLs co-located with those found in Cosenza et al. (2024): HD3, HD4, HD10, HD15, and HD17 had overlapping confidence intervals with QTLs found in the multi-parent population analysis; whereas QTLs overlapping with HD5, HD13 and HD22 were detected in single populations. For plant height, 14 of the 29 QTLs (Table 3) co-located with QTLs for the same trait in the multi-parent population. Most of them were detected in single populations, but three (PH15, PH22, PH24) were detected in the whole population. Regarding thousand grain weight, out of the 27 QTLs found in our study, 10 coincided with TGW QTLs in the multi-parent population (Shrestha et al., 2022).

The population studied by Shrestha et al. (2022) and Cosenza et al. (2024) is composed of two-row and six-row subpopulations and, thus, probably represents a larger genetic diversity space than our panel. Moreover, the multi-parent population they used is more amenable to the detection of small effect QTLs. In any case, the multiple shared locations for QTLs in our study and theirs is likely a result of the higher power of detection achieved with either approach, thus providing further confidence in the associations found in the present study. Therefore, the loci identified constitute a sound set of QTLs, with a clear potential to contribute to barley breeding in Europe.

A recent study (Hong et al., 2024), using a large and diverse two-rowed barley panel, found 38 MTAs for grain size, one in common with our TGW20 QTL. Finally, a meta-analysis revising yield-related traits of 54 studies that were published since 2000 (Du et al., 2024) found four QTLs co-locating with ours: TGW3, TGW17, TGW25 and GY2.

### Implications for breeding

The major selection footprints that we found appeared to be associated with disease resistance. We concentrated only on the most salient selective sweeps, although it is evident that breeding affected allelic frequencies throughout the genome with the shifts (large or small) paralleling always the time gradient (Fig. S10). The pericentromeric region of 5H, and its possible relation to the introduction of leaf rust tolerance, was thoroughly described in Wonneberger et al. (2023). Interestingly, the narrower, but equally marked selection footprint of 5HL appears to be associated with two stem rust resistance QTLs (Case et al., 2018). The selection footprint found in 4HL was not associated with agronomic QTLs. The closest one was TGW13, 15 Mb away. However, a QTL for net blotch tolerance, on the position of the selection footprints was found by Daba et al. (2020).

The dynamics of the evolution of allele frequencies for the agronomic QTLs in the timeframe of our study suggest an early selection of many plant height QTLs, followed closely in time by selection for thousand grain weight. Not surprisingly, both traits usually show high heritability. Effective selection for grain yield QTLs occurs later, apparently facilitated once the variance for the other traits was reduced. Plant height is easy to select, due to its high heritability, and shortness was preferred to minimize lodging, particularly after the introduction of semi-dwarf alleles like *denso*/*sdw1* (detected as PH14). Grain yield and thousand-grain weight were also important breeding targets for production and quality, but their selection appeared stronger after PH was improved. For instance, in the close QTL pairs PH1-TGW1, PH6-TGW4, PH15-TGW10, PH4-GY1, and PH8-GY3 (Table S4), PH-favourable alleles were fixed, or almost so, in the most modern class of cultivars, whereas the frequencies of favourable alleles for the QTLs of the other trait were low (although already growing in some cases). This was also the case for the region with QTL PH27-TGW26-GY9. In this region, modern varieties have favourable alleles for GY and PH almost fixed, while the favourable TGW allele frequency is still low. In other cases, favourable alleles for the two traits in a particular region were found in intermediate frequencies throughout the breeding history (PH10-TGW7, PH26-TGW25, PH13-TGW9, PH24-TGW21). Therefore, selection for both favourable alleles is likely feasible although, in the last two cases, the genotypic frequencies indicate a possible linkage in repulsion for the favourable alleles. In all other cases, the favourable alleles had already been selected. These observations can be of direct use for barley breeders.

GY7 showed the largest frequency shift, around 0.9, across time of all QTLs detected in this study. It was very effectively selected throughout the second half of the 20^th^ Century. The QTLs PH19 and TGW15 lie close to GY7. These two QTLs share the flag marker, although with opposite effects for agronomic fitness. One allele is associated with low height and thin grains, and the other one to tall plants and large grains. The antagonistic effect of a single gene on these two traits was already described for the main semi-dwarfing gene in barley, *HvGA20ox2* (Thomas et al., 1991). In this case, breeding selected short plants preferentially over large grains, although grain weight may have been compensated by selection at other loci. A possible explanation of the historic trends observed is that, once the PH19/TGW15 QTL was almost fixed for short plants, selection for grain yield at this region was facilitated. Indeed, favourable alleles for low TGW and plant height appear together in 142 cultivars, whereas the only nine cultivars with the “elevated height” allele also present the “high TGW” allele.

The near fixation of favourable alleles for traits which suffered strong selection pressure on neighbouring QTLs may have helped the selection at GY7. A similar situation may have occurred at 5HL, where the near fixation of the favourable (low height) allele at PH22, may have strengthened selection near TGW18, whose favourable allele suffered strong selection over time. The proximity of some QTLs and their fates throughout breeding suggests possible targets for future breeding. For example, in the distal region of 3HL, PH15 shows a decreasing frequency of “high” alleles, indicating selection pressure; however, the favourable allele of the neighbouring QTL, TGW10, remains at a low frequency. This suggests the presence of linked favourable alleles in repulsion, a situation that could be addressed by breeding. A similar phenomenon occurs at the beginning of 2H, with QTL PH6 and TGW4.

### Candidate genes

Some candidate loci corresponded to the already-known functions of characterised genes, while loci either provided new information about the effects of known genes or revealed genes with unknown roles in barley, but with suggested ones based on orthologues in rice (Table 5). For example, the candidate gene for HD17 is a homologue of *Arabidopsis thaliana AtNUP160*, associated with flowering time (Li et al., 2020). It anchors HOS1 to the ubiquitinated CONSTANS protein, which has not been reported in barley so far. Further evidence for the possible involvement of this gene in flowering comes from its preferential expression in barley apical meristems, inflorescences, and microspores (Li et al., 2023).

**Table 5.** Best candidate genes found for selected QTLs.

| Trait | QTL | CHR | Candidate gene | Gene model ID | MorexV3 Annotation |
| --- | --- | --- | --- | --- | --- |
| HD | HD1 | 1H | WNK3 | HORVU.MOREX.r3.1HG0004150 | Kinase family protein |
| HD | HD2 | 1H | HvCMF10 | HORVU.MOREX.r3.1HG0041150 | Zinc finger protein CONSTANS |
| HD | HD3 | 1H | HvCO9 | HORVU.MOREX.r3.1HG0058180 | CONSTANS-like protein |
| HD | HD5 | 2H | HvCO18 | HORVU.MOREX.r3.2HG0115000 | CONSTANS-like protein |
| HD | HD9 | 3H | HvFDL-H4 | HORVU.MOREX.r3.3HG0296180 | BZIP transcription factor |
| HD | HD16 | 5H | HvFRI | HORVU.MOREX.r3.5HG0534990 | FRIGIDA-like protein, putative |
| HD | HD17 | 6H | HvNUP160 | HORVU.MOREX.r3.6HG0557980 | Nuclear pore complex protein Nup 160 |
| HD | HD21 | 7H | HvMADS26 | HORVU.MOREX.r3.7HG0705340 | MADS-box transcription factor |
| HD | HD22 | 7H | HvNF-Yb2 | HORVU.MOREX.r3.7HG0734470 | Nuclear transcription factor Y subunit B |
| PH | PH1 | 1H | HvNRT | HORVU.MOREX.r3.1HG0005090 | Nitrate transporter 1.1 |
| PH | PH3 | 1H | OsCKX9 | HORVU.MOREX.r3.1HG0059670 | Cytokinin oxidase/dehydrogenase |
| PH | PH4 | 1H | HvGA20x8a | HORVU.MOREX.r3.1HG0087020 | Gibberellin 2-oxidase |
| PH | PH13 | 3H | OsWRKY21 | HORVU.MOREX.r3.3HG0297540<br>HORVU.MOREX.r3.3HG0297550 | WRKY transcription factor |
| PH | PH14 | 3H | HvGA20ox2 (sdw1/denso) | HORVU.MOREX.r3.3HG0307130 | Gibberellin 20 oxidase |
| PH | PH20 | 5H | OsPH9 | HORVU.MOREX.r3.5HG0496220 | Histone H4 |
| PH | PH24 | 6H | HvLAZY1 | HORVU.MOREX.r3.6HG0632820 | WRKY transcription factor-like protein |
| PH | PH29 | 7H | OsMYB45 | HORVU.MOREX.r3.7HG0736780<br>HORVU.MOREX.r3.7HG0736790<br>HORVU.MOREX.r3.7HG0736800<br>HORVU.MOREX.r3.7HG0736840 | Homeodomain-like superfamily protein<br>Homeodomain-like superfamily protein<br>Two-component response regulator<br>MYB transcription factor-like |
| TGW | TGW2 | 1H | OsCKX9 | HORVU.MOREX.r3.1HG0059670 | Cytokinin oxidase/dehydrogenase |
| TGW | TGW2 | 1H | HvSMOS1 | HORVU.MOREX.r3.1HG0058550 | AP2-like ethylene-responsive transcription factor |
| TGW | TGW7 | 3H | HvRA2 (vrs4) | HORVU.MOREX.r3.3HG0233930 | LOB domain protein |
| TGW | TGW8 | 3H | HvBRI1 | HORVU.MOREX.r3.3HG0285210 | Receptor kinase |
| TGW | TGW13 | 4H | HvGT1 | HORVU.MOREX.r3.4HG0399240 | Homeobox protein, putative |
| TGW | TGW14 | 4H | HvCMF4 | HORVU.MOREX.r3.4HG0411680 | Zinc finger protein CONSTANS |
| TGW | TGW18 | 5H | HvBRXL4 | HORVU.MOREX.r3.5HG0535760 | Protein BREVIS RADIX |
| TGW | TGW27 | 7H | HvJMJ | HORVU.MOREX.r3.7HG0752360 | Lysine-specific demethylase |
| GY | GY1 | 1H | HvGA20x8a | HORVU.MOREX.r3.1HG0087020 | Gibberellin 2-oxidase |
| GY | GY2 | 2H | HvHOX1 | HORVU.MOREX.r3.2HG0184740 | Homeobox leucine zipper protein |
| GY | GY5 | 3H | HvAGL2 | HORVU.MOREX.r3.3HG0221900 | Alpha-glucosidase |
| GY | GY8 | 7H | HvKAO1 | HORVU.MOREX.r3.7HG0637750 | Cytochrome P450 |
| GY | GY11 | 7H | OsMYB45 | HORVU.MOREX.r3.7HG0736780<br>HORVU.MOREX.r3.7HG0736790<br>HORVU.MOREX.r3.7HG0736800<br>HORVU.MOREX.r3.7HG0736840 | Homeodomain-like superfamily protein<br>Homeodomain-like superfamily protein<br>Two-component response regulator<br>MYB transcription factor-like |

Looking at the confidence intervals of HD QTLs, several known flowering-related genes were located within them. Among these 22 QTLs, the confidence regions included *HvCO9* (Cockram et al., 2012), *HvFT3* (Faure et al., 2007, Kikuchi et al., 2009), *HvHAP3* (Campoli et al., 2013), and *HvVRN2* (Karsai et al., 2005), respectively in HD3, HD4, HD6, and HD13. However, *HvFT3* is not a good candidate, since all 151 cultivars have the same (presence) allele at this gene. Similarly, *HvVRN2* is not a likely candidate, as the allelic segregation for this gene does not coincide with that of the QTL (Table S1). On the contrary, three more QTLs were located near flowering-related genes that are good candidates: HD13 was only 3 Mb apart from *HvFT5* (Faure et al., 2007, Kikuchi et al., 2009), HD21 was 5.5 Mb away from *HvMADS26* (Pankin et al., 2018, Hill et al., 2019), and HD16 was 383 kb away of *HvFRI* (Campoli et al., 2013).

The candidate gene for the plant height QTL PH24 corresponds to the orthologous one in rice, *LAI1/LAZY1*. This gene regulates the expression of auxin transporters to control tiller angle and shoot gravitropism (Li et al., 2007; Zhu et al., 2020). Interestingly, a candidate gene for the thousand-grain weight QTL TGW18 is the orthologue of rice *OsBRXL4,* which is a regulator of the nuclear localization gene of rice *LAI1/LAZY1* (Li et al., 2019). Although neither rice gene has been studied for grain weight, they are highly related to plant architecture. There is a possible candidate locus affecting two QTLs on 7HL, PH29 and GY11. The candidate is a cluster of four orthologues of the rice gene *OsMPH1*/*OsMYB45*, located exactly in the overlapping interval of the two QTLs. This rice gene affects plant height and grain yield (Zhang et al., 2017), which is consistent with our observations. The favourable alleles at these two QTL were present at high frequencies, and both increased over time until fixation, indicating joint selection in this region.

The *HvHOX1* (*Vrs1)* gene is a candidate for QTL GY2. The haplotype *Vrs1.t*, commonly known as *deficiens* (Sakuma et al., 2017) could underlie the favourable allele, providing a yield advantage of 0.28 t ha^-1^ across all trials. Selection at this QTL apparently started in the 1980-90s and was subject to strong selection pressure. This allele conferred larger grains, although this did not result in a yield advantage in the study by Sakuma et al (2017). Those authors hypothesized that *deficiens* induces larger grains by the suppression of organs (lateral florets), not specifically involved in sink/source relationships. In our study, however, this QTL did not show a clear signal for TGW, but the GY response was very consistent across environments. To the best of our knowledge, this is the first time that a yield advantage has been reported for this gene under field conditions.

In summary, our results indicate when and where breeding removed allelic diversity, which traits were more accessible to breeders, where there were effects on nearby genes, and how. When combined with candidate gene identification, our approach allows the biological function of gene under relevant field conditions to be examined. This information, in sum, provides a road map to help breeders fine tuning their targets for the future.

## Supplementary data

Supplementary Table S1. List of 164 genotypes, with country and release year. Allelic data for flowering-related genes and *Vrs1*.

Supplementary Table S2. Information of the field trial network and best spatial correction models.

Supplementary Table S3. Summary of MTAs of single-trial GWAS.

Supplementary Table S4. Co-localisation of meta-QTLs between traits.

Supplementary Table S5. Summary of meta-QTLs enrichment with exome capture markers.

Supplementary Table S6. BLUEs of the four phenotypic traits evaluated.

Supplementary Table S7. Genotypic data of 151 two-rowed spring barleys

Supplementary Figure S1. PCA of the 164 genotypes, showing which were selected for further analyses.

Supplementary Figure S2. Correlation between trials for each trait.

Supplementary Figure S3. Kinship matrix of the selected 151 genotypes.

Supplementary Figure S4. Boxplots for all phenotypes and trials, divided by groups of release year.

Supplementary Figure S5. Linear relationships between grain yield with plant height and thousand-grain weight.

Supplementary Figure S6. Biplots of AMMI analyses for plant height, thousand-grain weight and grain yield.

Supplementary Figure S7. Differences in heading time between the mean of each trial with genotypes Saana and Mona.

Supplementary Figure S8. Manhattan plots of the meta-GWAS, for flowering time, plant height and thousand-grain weight.

Supplementary Figure S9. Changes in allele frequencies over time of flowering time and thousand-grain weight meta-QTLs.

Supplementary Figure S10. Genome-wide fingerprints of selection.

Supplementary Figure S11. Photo of the spikes from the 14 genotypes with the favourable allele of QTL GY2, linked to *Vrs1*.

## Acknowledgements

The authors are grateful to all technicians who participated in both European projects.

## Author contributions

Funding was acquired by AMC, LC, BK, AG, RW, AJF, KP, AHS, LR, EI. The study was conceptualized by AMC, AT, LC, BK, JR, RW, WT, KP, AHS, AV, LR and EI. Resources were provided by AMC, LC, BK, AG, RW, KP, AHS, LR. Field experiments and data collection were carried out by AMC, AT, SD, RS, CC, JR, LR, WT, FS, KP, AHS, MJ, FLS, MP, AV, SS and EI. Methodology was designed by FMT, AT, LR, WT, SS, EI. Data curation was carried out by FMT, AT, WT, AJF, SS and EI. Analyses were performed by FMT, AT, WT and EI. The study was managed by AMC, LC, BK, RW, WT, KP, AHS, FLS, AV, LR. Analyses and plotting code were written by FMT, WT and EI. The research was supervised AMC, AT, LC, BK, RW, WT, KP, AHS, FLS, AV and LR. Plots were created by FMT, AMC and EI. Original draft was prepared by FMT, AMC, AT and EI. All authors revised the final manuscript.

## Conflict of interest

The authors declare no conflict of interest.

## Funding

This work was supported by the ERA-PG-funded project Exbardiv (Genomics-Assisted Analysis and Exploitation of Barley Diversity; http://www.erapg.org), the FACCE ERA-NET Plus project ClimBar 618105 (An integrated approach to evaluate and utilise genetic diversity for breeding climate-resilient barley), the SUSCROP ERA-NET project BARISTA PCI2019-103758 (Advanced tools for breeding BARley and Intensive and SusTainable Agriculture under climate change scenarios), and by the Government of Aragón grants A08_20R and A08_23R. FMT PhD scholarship was funded by the Government of Aragón, 2019-2023. FMT one-month traineeship at James Hutton Institute, under the supervision of LR, was funded by CSIC mobility project LINKB20052 (Barley adaptation under climate change. Timing and efficiency of flowering).

## Data availability

The phenotypic and genotypic data that support the findings of this study are provided as supplementary Tables S6 and S7, respectively.

## Notes

### Competing Interest Statement

The authors have declared no competing interest.

